# Epstein-Barr Virus Latent Membrane Protein 1 targets cIAP1, cIAP2 and TRAF2 for Proteasomal Degradation to Activate the Non-canonical NF-κB Pathway

**DOI:** 10.1101/2025.09.06.674652

**Authors:** Yizhe Sun, Shunji Li, Bidisha Mitra, Ling Zhong, Aretina Zhang, Benjamin E. Gewurz

## Abstract

The Epstein-Barr virus oncoprotein Latent Membrane Protein 1 (LMP1) is expressed in multiple malignancies and is critical for B-cell immortalization. LMP1 constitutively activates NF-κB signaling pathways, which are essential for EBV-mediated B cell transformation and for transformed B cell survival. Reverse genetic analysis revealed two LMP1 regions critical for primary human B cell immortalization, termed transformation effector site (TES) 1 and 2, which activate multiple host growth and survival pathways, in particular NF-κB. Of these, only TES1 signaling is required for B-cell transformation within the first several weeks of infection. TES1 signaling is also critical for EBV-transformed lymphoblastoid B-cell survival. However, precisely how TES1 initiates NF-κB signaling has remained incompletely understood. Here, we provide multiple lines of evidence that TES1 associates with cellular inhibitor of apoptosis protein 1 and 2 (cIAP1/2) in a tumor necrosis factor associated factor 3 (TRAF3) dependent manner. TES1 signaling drives cIAP1 autoubiquitination and targets TRAF2, cIAP1 and 2 for proteasomal degradation in a TRAF3 dependent manner. Overexpression of either cIAP1 or 2 impaired LMP1 TES1-mediated non-canonical NF-κB activation. Collectively, these studies suggest that LMP1 TES1 initiates non-canonical NF-κB signaling distinctly from CD40 and other host immunoreceptors, thereby highlighting a therapeutic target.

## Introduction

The gamma-herpesvirus Epstein-Barr virus (EBV) establishes lifelong infection of nearly 95% of adults worldwide. EBV causes ∼200,000 cancer cases per year, including multiple types of lymphomas, gastric and nasopharyngeal carcinoma[1-4]. These include Burkitt lymphoma, Hodgkin lymphoma, primary central nervous system (CNS) lymphoma, post-transplant lymphoproliferative disease (PTLD), T and NK cell lymphoma[5]. EBV is also a major trigger of autoimmune diseases, in particular multiple sclerosis and systemic lupus erythematosus[6-9].

EBV subverts key host pathways to drive proliferation and differentiation of newly-infected B cells into memory cells, the reservoir for lifelong infection. To do so, EBV expresses a series of latency oncogenes, comprised of six Epstein-Barr nuclear antigens (EBNA) and two latent membrane proteins (LMP), together with non-coding RNAs (ncRNA). Latency programs utilize distinct combinations of viral latency genes[10-13]. The fully transforming latency III program, comprised of all six EBNA, LMP1, LMP2A, LMP2B and ncRNAs drives B-cells into secondary lymphoid germinal centers. If left unchecked by immune pressure, latency III immortalizes infected cells into lymphoblastoid B cell lines (LCLs). Latency III cells are observed in PTLD and primary CNS lymphoma. Within germinal centers, infected B cells switch to the latency II program, where EBNA1, LMP1 and 2A are expressed with ncRNA. Latency II is observed in Hodgkin lymphoma Reed-Sternberg tumor cells, in T and NK lymphomas, and also in a subset of nasopharyngeal carcinoma[14, 15]. Upon memory cell differentiation, EBV switches to latency I, where EBNA1 is the only viral protein-coding gene expressed.

LMP1 expression is sufficient to drive rodent fibroblast transformation and polyclonal B-cell proliferation[16-18]. LMP1 is comprised of a 24-residue cytoplasmic N-terminal tail, six transmembrane domains and a 200 residue C-terminal cytoplasmic tail[11, 13, 19]. LMP1 transmembrane domains drive lipid raft association and continuous, ligand independent signaling from C-terminal tail regions[20-23]. Of these, reverse genetic studies defined that signaling from transformation effector sites (TES) 1 and 2, also referred to as C-terminal activating regions (CTAR) 1 and 2, are necessary for EBV-mediated primary B-cell transformation[20, 24].

TES1/CTAR1 spans residues 186-231 and uses a PXQXT motif to engage tumor necrosis factor receptor associated factors (TRAFs) to drive downstream non-canonical NF-κB, MAP kinase and PI3K pathways[25-29]. TRAF1 enables TES1 to also potently activate canonical NF-κB[28]. TES2/CTAR2 residues 351-386 recruits TRAF6[30] independently triggers canonical NF-κB, MAPK, IRF7, and P62 pathways[31, 32]. The LMP1 C-terminal tail CTAR3 region also activates JAK/STAT and SUMOylation pathways[33]. Of these, only TES1 signaling is necessary for EBV B cell transformation[20, 24-29, 34, 35].

LMP1 mimics aspects of signaling by the B-cell co-receptor CD40, whose binding by CD40 ligand stimulates NF-κB, MAP kinase activity. CD40 signaling is required for B-cell development, germinal center formation, class switch recombination and somatic hypermutation[36, 37]. Transgenic mice that express the LMP1 C-terminal tail in place of that of CD40 have normal B-cell development, activation and immune responses, though do exhibit cytokine-independent class-switch recombination[38]. Similarly, mice with transgenic B-cell LMP1 or chimeric LMP1/CD40 cytoplasmic tail have largely overlapping phenotypes[39]. However, since LMP1 signals constitutively and in a ligand-independent manner, its expression is transforming. Transgenic LMP1 expression transforms rodent fibroblasts[16]. In the absence of T and NK surveillance, transgenic LMP1 expression drives fatal B-cell proliferation, particularly when co-expressed with LMP2A, which mimics aspects of B-cell immunoglobulin signaling[12, 40, 41]. LMP1 and EBNA2 co-expression are sufficient to transform human peripheral blood B cells into immortalized lymphoblasts[42], further suggesting LMP1 as a major EBV therapeutic target.

The non-canonical NF-κB pathway promotes B cell survival, maturation and homeostasis in peripheral lymphoid organs, and has key roles in germinal center formation[43, 44]. Impairment of non-canonical NF-κB signaling results in immunodeficiency, whereas its hyperactivation instead contributes to autoimmune, inflammatory and neoplastic diseases[43]. Non-canonical NF-κB is suppressed at baseline by constitutive targeting of the NF-κB activating kinase (NIK) for ubiquitin-mediated proteasomal degradation. A complex containing TRAFs 2, 3, cellular inhibitor of apoptosis (cIAP) 1 and 2 binds to NIK, typically using TRAF3 as an adaptor protein[43, 44]. The E3 ubiquitin ligase activity of cIAP 1 and 2 targets NIK for proteasomal degradation to suppress non-canonical NF-κB signaling in the absence of stimuli.

Through multiple distinct mechanisms, non-canonical pathways disrupt cIAP-mediated NIK turnover to activate downstream signaling. For instance, to initiate signaling, CD40 and B-cell activating factor (BAFF) receptors trigger cIAPs to ubiquitinate and slate TRAF3 for proteasomal degradation, thereby preventing NIK ubiquitination by the TRAF2/cIAP1/cIAP2 complex. To do so, they increase K63-linked polyubiquitin chain attachment to TRAF2[45, 46]. Increased NIK abundance results in its autophosphorylation, which stimulates its kinase activity towards the kinase IKKα. Activated IKKα phosphorylates p100, which triggers its partial proteasomal processing into the active p52 NF-κB transcription factor subunit. p52 traffics into the nucleus as homodimers or heterodimers, including with RelB, to activate non-canonical pathway targets[43]. Comparatively less is known about how TES1 initiates non-canonical NF-κB signaling.

The LMP1 PXQXT motif acts as a docking site for TRAFs 1, 2, 3 and 5[34, 47-49]. A substantial amount of the LCL TRAF1 and 3 pools are associated with LMP1, whereas a comparatively low amount of TRAF2 is LMP1 associated[26, 50-52]. TRAF3 binds more tightly to LMP1 than other TRAFs and can compete with TRAF1 and TRAF2 for LMP1 association[26, 50, 51]. It has been suggested that TES1 sequesters TRAF3 to initiate non-canonical NF-κB signaling[53]. LMP1 signaling was intact in TRAF2-/- B cells[48, 54], though TRAF2 knockout itself activates non-canonical NF-κB, complicating these analyses. TES1 signaling activates non-canonical NF-κB through NIK and IKKα[27, 55-59].

A range of LMP1 target genes have been defined[60, 61]. Key TES1 targets include the anti-apoptotic factor cFLIP, which is necessary for LCL survival[29, 62], underscoring the importance of TES1 signaling. Yet, the mechanism by which LMP1 initiates non-canonical NF-κB signaling has remained incompletely defined. Interestingly, despite its central role in control of other non-canonical pathways, cIAP activity has not been investigated in the context of LMP1 signaling. Here, we report that LMP1 associates with both cIAP1 and 2, in a TRAF-dependent manner. We present evidence that LMP1 recruits and targets cIAP1, 2 and TRAF2 for proteasomal degradation in a TRAF3-dependent manner to initiate non-canonical NF-κB signaling, suggesting a previously unappreciated manner by which LMP1 TES1 drives this key oncogenic pathway.

## Results

### LMP1 TES1 signaling downmodulates TRAF2, cIAP1 and cIAP2 abundances

To gain insights into how LMP1 initiates non-canonical NF-κB signaling, we used EBV-negative Daudi and Akata Burkitt B-cell lines with doxycycline inducible expression of wildtype (WT) vs well-characterized LMP1 TES1 vs TES2 mutant alleles, in which point mutations abrogate signaling from TES1 and/or TES2[29, 63, 64]. The TES1 mutant (TES1m) harbored two alanine point mutations in the TRAF binding motif (204PQQAT208 → AQAAT) that impede TRAF recruitment, whereas the TES2 mutant (TES2m) 384YYD386→ID substitutions block TES2 signaling. We also utilized an LMP1 TES1 and TES2 double mutant (DM) with both of these mutations (**Fig. 1A**). We validated that each had similar levels of LMP1 expression at 24 hours post-induction by 250 ng/ml doxycycline, that LMP1 with signaling from either or both TES induced TRAF1 albeit to varying degrees as expected[29], and that non-canonical NF-κB activation was highly impaired by TES1m or DM LMP1, as expected (**Fig. 1B**).

**Figure 1.**
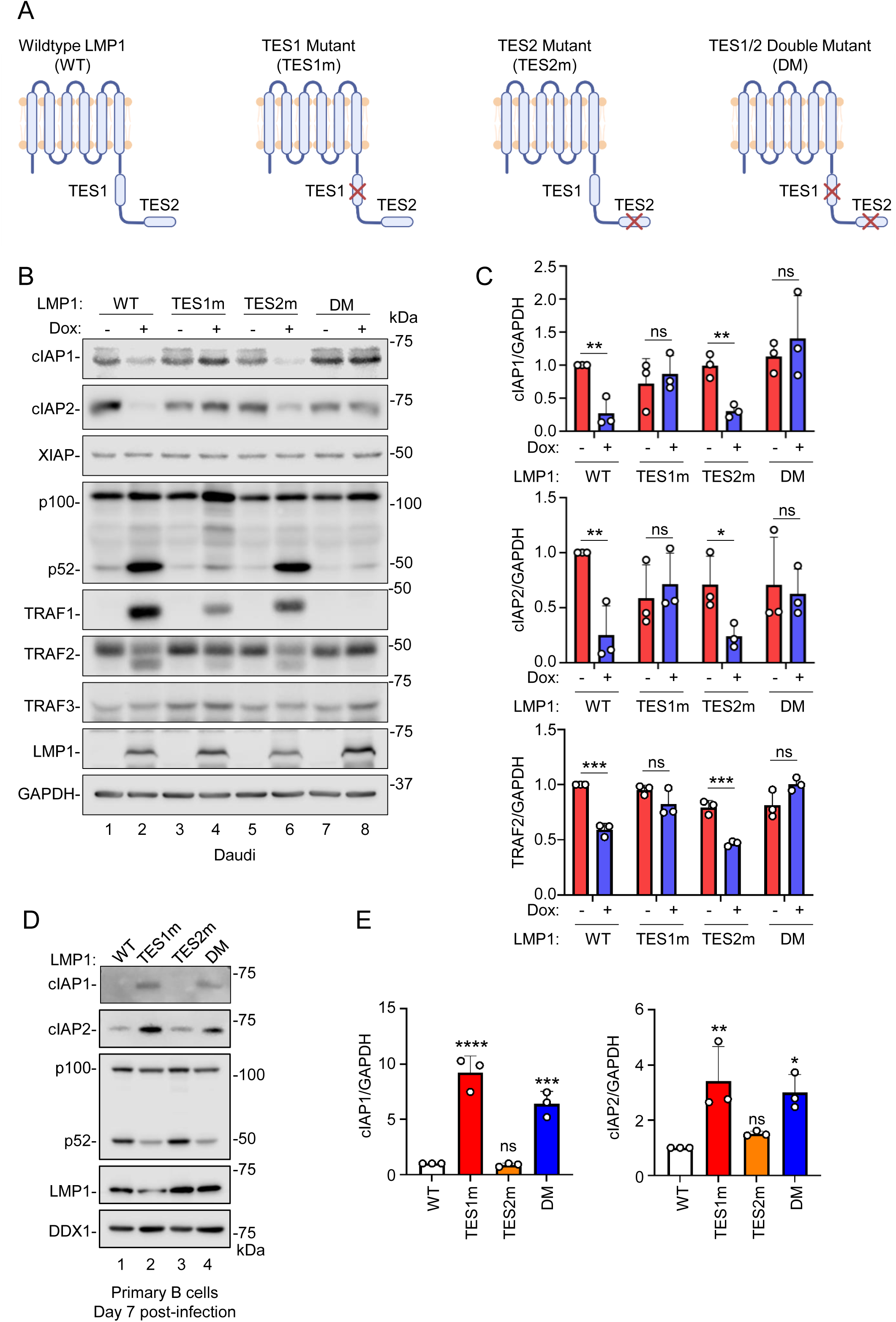
LMP1 TES1 signaling downregulates cIAP1, cIAP2 and TRAF2 expression. (A) Schematic diagram of wildtype (WT), TES1 point mutant (TES1m), TES2 point mutant (TES2m), or TES1/2 double mutant (DM) LMP1. (B) Analysis of LMP1 effects on cIAP1, cIAP2, XIAP, and TRAFs expression in Daudi Burkitt cells. Immunoblot analysis of whole cell lysates (WCL) from Cas9+ Daudi Burkitt B-cells induced for WT, TES1m, TES2m, or DM LMP1 expression by addition of 250ng/mL doxycycline (Dox) for 24 hours. (C) Relative fold changes + standard deviation (SD) of GAPDH load control normalized cIAP1, cIAP2 and TRAF2 levels, based on densitometry from three replicates as in (B). Values in vehicle control treated WT LMP1 expressing cells were set to 1. (D) Analysis of LMP1 effects on cIAP1 and cIAP2 expression in newly infected primary B-cells. Immunoblot analysis of WCL from human peripheral blood B cells infected with EBV expressing WT, TES1m, TES2m or DM LMP1 at day 7 post-infection. DDX1 was used as a load control, since its protein levels do not change significantly following primary B cell infection by EBV[96, 97]. (E) Relative fold changes + SD of GAPDH load control normalized cIAP1 and cIAP2 levels, based on densitometry from three replicates as in (D). Values in vehicle control treated WT LMP1 expressing cells were set to 1. Statistical significance was assessed by two-tailed unpaired Student’s t-test (C) or one-way ANOVA followed by Tukey’s multiple comparisons test (E). ns, not significant, *p<0.05, **p<0.01, ***p<0.001, ****p<0.0001. Blots (B and D) are representative of n=3 experiments.

While cIAP ubiquitin ligases have major roles in control of most non-canonical NF-κB pathways[43], they have not yet been characterized downstream of LMP1. We were therefore intrigued to find that cIAP1 and 2 were highly depleted in both Daudi and Akata B cells 24 hours after induction of expression of WT or TES2m LMP1, in which TES1 signaling was active. By contrast, we did not observe significantly changed cIAP1 or 2 expression following induction of similar levels of TES2m or DM LMP1, in which TES1 signaling was inactive (**Fig 1B-C and S1**). Notably, levels of inhibitor of apoptosis family member XIAP remained unchanged upon expression of any of the LMP1 constructs (**Fig 1B and S1A**), suggesting a potentially specific TES1 effect at the level of cIAP1 and 2. We also noticed that induction of WT or TES2m LMP1 resulted in diminished levels of TRAF2, but not TRAF3 (**Fig 1B-C and S1**). Furthermore, we noted that a lower molecular weight band immunoreactive with anti-TRAF2 antibody appeared following induction of WT or TES2m expression (**Fig 1B and S1A**), suggesting that TES1 signaling may induce TRAF2 cleavage. We note that CD40 signaling induces TRAF2, but not cIAP1/2 degradation[65, 66].

As a complimentary approach, we assessed cIAP1 and 2 levels in primary human B-cells infected by recombinant EBV with wildtype, TES1m, TES2m or DM LMP1. Of note, LMP1 is not required for the first 8 days of newly infected primary B-cell outgrowth[67]. We observed significantly lower cIAP1 and cIAP2 levels in cells infected by EBV encoding WT or TES2m LMP1, in which TES1-mediated non-canonical NF-κB signaling was active, as evidenced by robust p100:p52 processing (**Fig. 1D-E**). Taken together, these results raise the possibility that TES1 signaling may target cIAP1 and cIAP2 for degradation.

### Latency III destabilizes TRAF2, cIAP1 and cIAP2 in B-cells

Constitutive LMP1/NF-κB signaling in latency III cells drives synthesis of both cIAP1 and cIAP2, which are well-characterized NF-kB target genes[68]. In contrast to the loss of cIAP1/2 and TRAF2 seen upon induction of conditional LMP1 expression in Burkitt cells, expression of each is detectable in latency III Burkitt cells and in LCLs. We therefore hypothesized that their steady state levels in latency III cells represent the balance of their synthesis and turnover. To test this, we performed cycloheximide (CHX) chase analysis, using isogenic MUTU Burkitt cells with latency I vs III programs, termed MUTU I vs III[69]. LMP1 displayed a half-life of 4 hours, consistent with previous research[70]. Interestingly, cIAP1 and cIAP2 exhibited shorter half-lives in MUTU III than in MUTU I, with T_1/2_ < 1 hour in MUTU III but > 4 hours in MUTU I (**Fig. 2A-B**). Similarly, TRAF2 levels decreased by ∼50% over the first four hours of cycloheximide chase in MUTU III, but were little changed in MUTU I at this time point (**Fig. 2A-B**). Intriguingly, we observed two TRAF2 species in immunoblots of MUTU III whole cell lysates (WCL), which migrated at a similar molecular weight and at a somewhat lower molecular weight than in blots of MUTU I WCL (**Fig. 2A**). These data are consistent with a model in which LMP1 targets cIAP1/2 for degradation, but also induces cIAP1/2 expression through NF-κB target gene regulation, potentially in a manner dependent on TRAF2 proteolysis.

**Figure 2.**
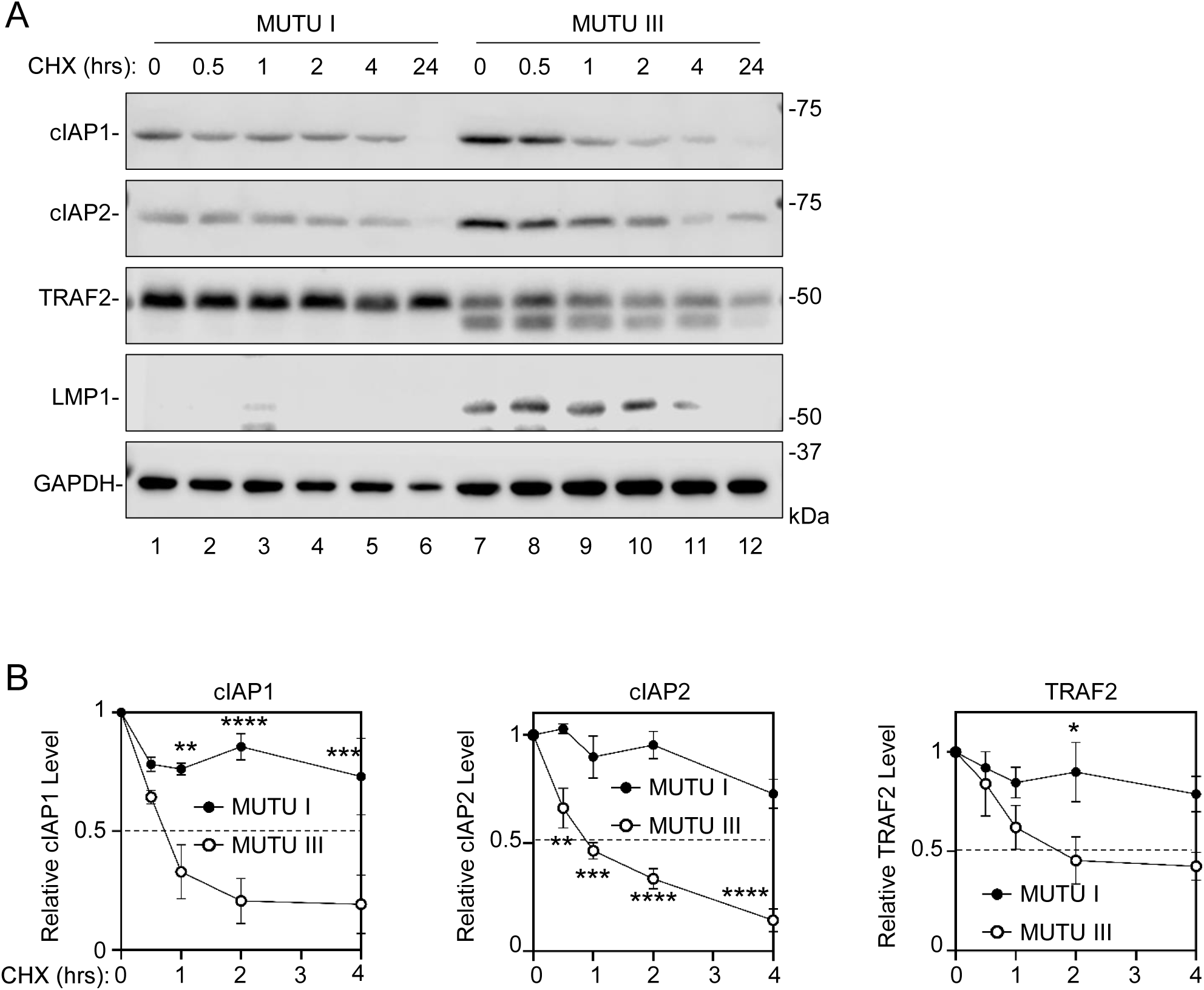
EBV latency III destabilizes cIAP1, cIAP2 and TRAF2. (A) Analysis of cIAP1, cIAP2, and TRAF2 half-lives in isogenic MUTU Burkitt B-cells that differ only by EBV latency I vs III programs. Shown are representative immunoblot analyses of WCL from MUTU I vs MUTU III cells treated with 50 µg/mL cycloheximide (CHX) for the indicated number of hours. Blots are representative of n=3 experiments. (B) Relative fold changes + SD of GAPDH load control normalized cIAP1, cIAP2 and TRAF2 levels, based on densitometry from three replicates as in (A). Values in MUTU I and III cells at 0 hours of CHX chase were set to 1. Statistical significance was assessed by two-way ANOVA followed by Tukey’s multiple comparisons test (B). ns, not significant, *p<0.05, **p<0.01, ***p<0.001, ****p<0.0001.

### TRAF3 is required for LMP1-mediated cIAP1/2 degradation

Given that TES1 recruits multiple TRAFs[26], and that particular TRAFs can recruit cIAPs to immunoreceptor signaling regions[71, 72], we next asked whether specific TRAFs were necessary for TES1 targeting of cIAP1/2. We therefore used CRISPR-Cas9 editing to selectively deplete either TRAF1, 2, 3 or 5, each of which associate with the TES1 PxQxT motif[26, 50, 51, 73-76]. Following single guide RNA (sgRNA) expression in Cas9+ Daudi Burkitt B-cells, immunoblot was used to confirm successful on-target TRAF depletion. We also confirmed that TRAF3 depletion was sufficient to trigger non-canonical NF-κB activity, as judged by p100/p52 processing (**Fig. 3A**). Intriguingly, depletion of TRAF3, but not of TRAFs 1, 2 or 5, significantly stabilized cIAP1 and cIAP2 upon conditional WT LMP1 expression (**Fig. 3A-B, S2**). As expected, TES1m LMP1 expression failed to deplete cIAP1 or 2 in cells depleted of TRAF1,2,3 or 5 (**Fig. 3A-B, S2**). Since TRAF2 and 5 have partially redundant function[77-80], it was however notable that conditional WT, but not TES1m LMP1 expression, also partially depleted TRAF5 (**Fig. S2C**).

**Figure 3.**
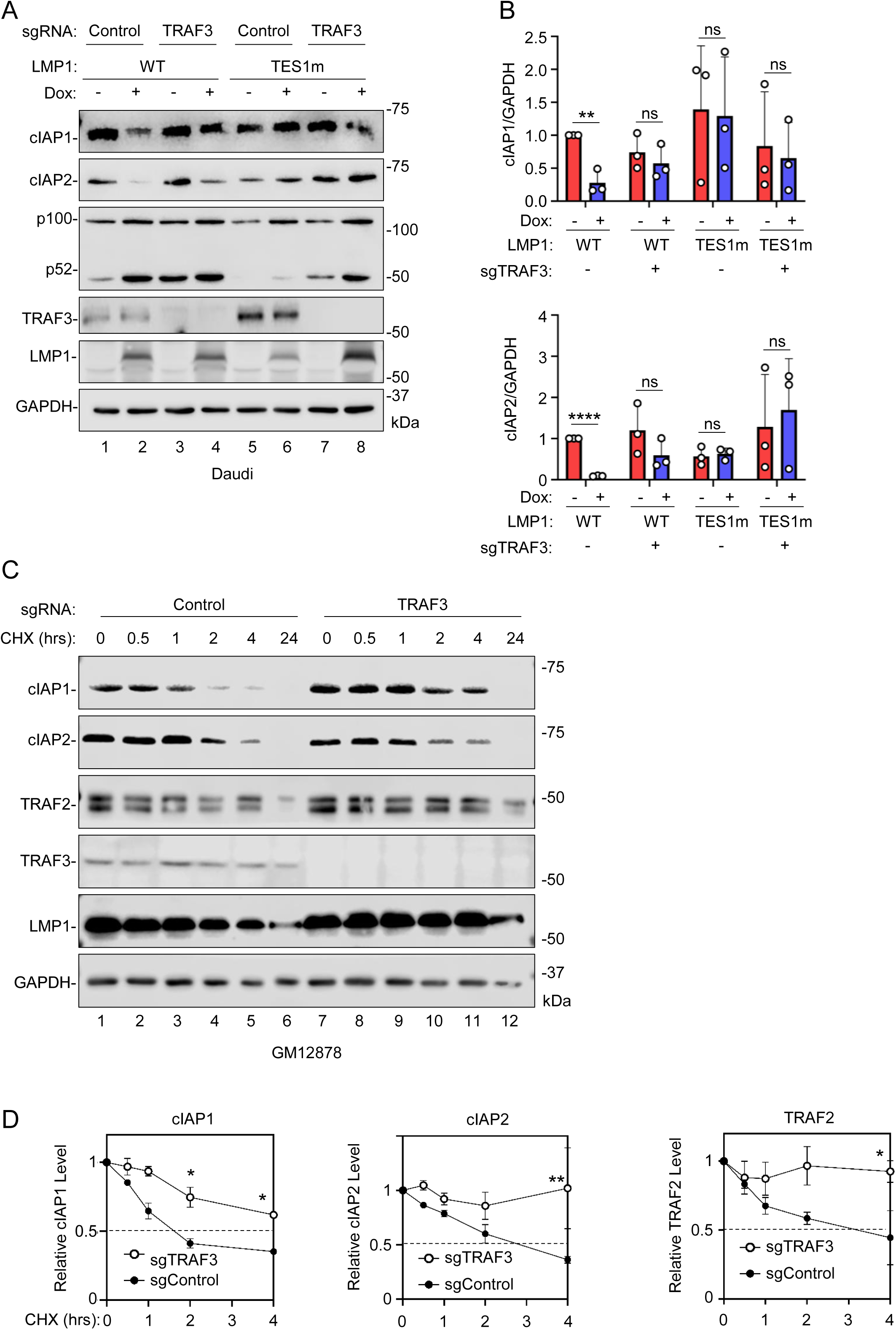
TRAF3 supports LMP1-induced cIAP1/2 depletion. (A) Analysis of TRAF3 roles in LMP1 TES1-mediated cIAP1 and cIAP2 depletion. Immunoblot analysis of WCL from Cas9+ Daudi cells that expressed control vs TRAF3 targeting single guide RNA (sgRNA) and that were induced for LMP1 expression by Dox (250 ng/mL) for 24 hours. (B) Relative fold changes + SD of GAPDH load control normalized cIAP1, cIAP2, based on densitometry from three immunoblot replicates as shown in (A). Values in Daudi cells with control sgRNA and uninduced for WT LMP1 expression were set to 1. (C) Analysis of cIAP1, cIAP2, and TRAF2 half-lives in control vs TRAF3 knockout (KO) GM12878 LCLs. Immunoblot analysis of WCL from Cas9+ GM12878 LCLs that expressed control versus TRAF3 targeting sgRNA and that were treated with 50µg/mL CHX for the indicated hours (hrs). (D) Relative fold changes + SD of GAPDH load control normalized cIAP1, cIAP2 and TRAF2 levels, based on densitometry from three replicates as in (C). Values in GM12878 cells with control sgRNA expression at 0 hours of CHX chase were set to 1. Statistical significance was assessed by two-tailed unpaired Student’s t-test (B) or by two-way ANOVA followed by Tukey’s multiple comparisons test (D). ns, not significant, *p<0.05, **p<0.01, ****p<0.0001. Blots are representative of n=3 experiments.

To further assess TRAF3 roles in cIAP1/2 turnover in the latency III context, we next performed cycloheximide chase analysis in the well-characterized LCL GM12878. CRISPR TRAF3 depletion again stabilized cIAP1 (**Fig. 3C**), even though TRAF3 itself exhibited remarkable stability over the 24-hour chase in control cells. TRAF3 depletion also prolonged TRAF2 half-life (**Fig. 3C-D**), suggesting that TES1 signaling may target a complex containing TRAF2, potentially also TRAF5, cIAP1 and cIAP2 for degradation in a TRAF3 dependent manner.

### LMP1 association with cIAP1/2 is dependent on TRAF3, but not TRAFs1, 2 or 5

We next examined whether LMP1 associated with cIAP1/2 in B-cells in a TES1-dependent manner. C-terminally HA-tagged LMP1 co-immunoprecipitated TRAFs 2 and 3, as expected, but also co-immunoprecipitated cIAP1 and cIAP2 (**Fig. 4A**). Importantly, TES1m and DM LMP1 failed to co-immunoprecipitate TRAFs 2/3 or cIAP1/2, suggesting that their association with LMP1 was dependent on the TES1 PxQxT motif (**Fig. 4A**). Similarly, confocal immunofluorescence analysis demonstrated LMP1 and cIAP1 colocalization, which was again dependent on an intact TES1 PxQxT sequence (**Fig. S3**).

**Figure 4.**
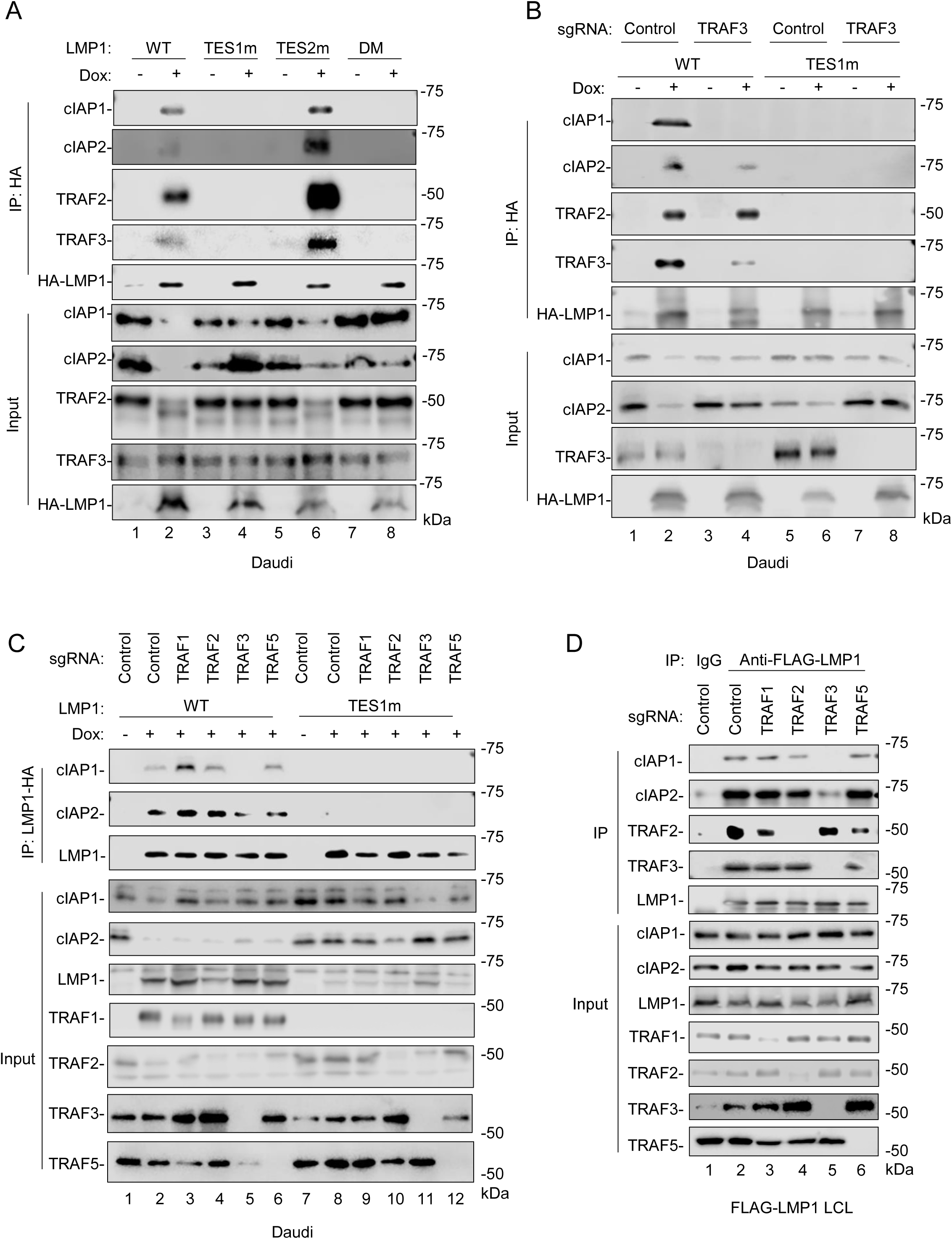
TRAF3 supports LMP1 association with cIAP1 and 2 in a TES1-dependent manner. (A) Analysis of LMP1 TES1 versus TES2 association with cIAP1 and cIAP2. Immunoblot analyses of 1% input vs anti-HA-LMP1 complexes immunopurified from Daudi cells mock induced or induced for HA-tagged WT, TES1m, TES2m or DM LMP1 for 24 hours by 250 ng/ml Dox. (B) Analysis of TRAF3 role in LMP1 association with cIAP1/2. Immunoblot analyses of 1% input vs anti-HA-LMP1 complexes immunopurified from Cas9+ Daudi cells that expressed control versus TRAF3 targeting sgRNA and induced for WT or TES1m LMP1 expression by 250ng/mL Dox for 24 hours. (C) Analysis of TRAF1, 2 and 5 roles in LMP1 association with cIAP1/2. Immunoblot analyses of 1% input vs anti-HA-LMP1 complexes immunopurified from Cas9+ Daudi cells that expressed control versus TRAF1,2,3 or 5 targeting sgRNA and induced for WT or TES1m LMP1 expression by 250ng/mL Dox for 24 hours. (D) Analysis of LCL TRAF roles in LMP1 association with cIAP1/2. Immunoblot analyses of 1% input vs anti-FLAG-LMP1 complexes immunopurified from Cas9+ LCLs that express FLAG-tagged LMP1 at physiological levels and that also expressed control versus TRAF1,2,3 or 5 targeting sgRNA. Anti-mouse IgG was used as a negative control. Blots are representative of n=3 (A and B) or 2 (C and D) experiments.

As TRAFs recruit cIAP1/2 to receptors[44, 45, 71, 81], we next examined whether specific TRAFs were necessary for cIAP1/2 association with LMP1. CRISPR TRAF3 depletion strongly diminished cIAP1 association and to a somewhat lesser extent cIAP2 association with HA-LMP1 in Daudi cell extracts (**Fig. 4B**). Importantly, TRAF3 depletion did not impair recruitment of TRAF2, which associated with LMP1 in a TES1 PxQxT motif-dependent fashion (**Fig. 4B**). By contrast, CRISPR depletion of either TRAFs 1, 2 or 5 did not substantially diminish cIAP1 or cIAP2 co-immunoprecipitation with HA-LMP1 (**Fig. 4C**). We extended this result into the latency III setting, using Cas9+ LCLs that express an N-terminally FLAG-tagged LMP1 allele knocked into the EBV genome[50]. FLAG-LMP1 co-immunoprecipitated cIAP1, 2 and TRAF2 in a manner dependent on TRAF3, but not TRAFs1, 2 or 5 (**Fig. 4D**). By contrast, TRAF2 associated with LMP1 in control LCLs or in LCLs depleted for TRAFs 1, 3 or 5, suggesting specificity of the cIAP1/2 result. Altogether, these data suggest that LMP1 TES1 associates with TRAF3-containing TRAF heterotrimers, which likely also include TRAF2 and which in turn recruit cIAP1 and cIAP2.

### LMP1 TES1 triggers cIAP1 polyubiquitination and cIAP1/2 proteasomal degradation

To gain insights into how TES1 signaling results in TRAF2 and cIAP1/2 turnover, we performed immunoblot analysis on whole cell lysates from Daudi cells treated with vehicle, the proteasome inhibitor MG132 or with the lysosome acidification inhibitor NH₄Cl followed by induction of WT or TES1m LMP1. TES1-dependent depletion of cIAP1 and cIAP2 was evident in vehicle control and NH₄Cl-treated cells, but to a considerably lesser degree in MG132-treated cells (**Fig. 5A-B**). MG132 mildly stabilized TRAF2, with the majority of TRAF2 species shifting to a lower molecular weight in control and MG132-treated Daudi cells induced for WT LMP1, but not for TES1m LMP1 expression (**Fig. S4**), suggesting that partial proteasomal digestion does not result in the modified TRAF2 species.

**Figure 5.**
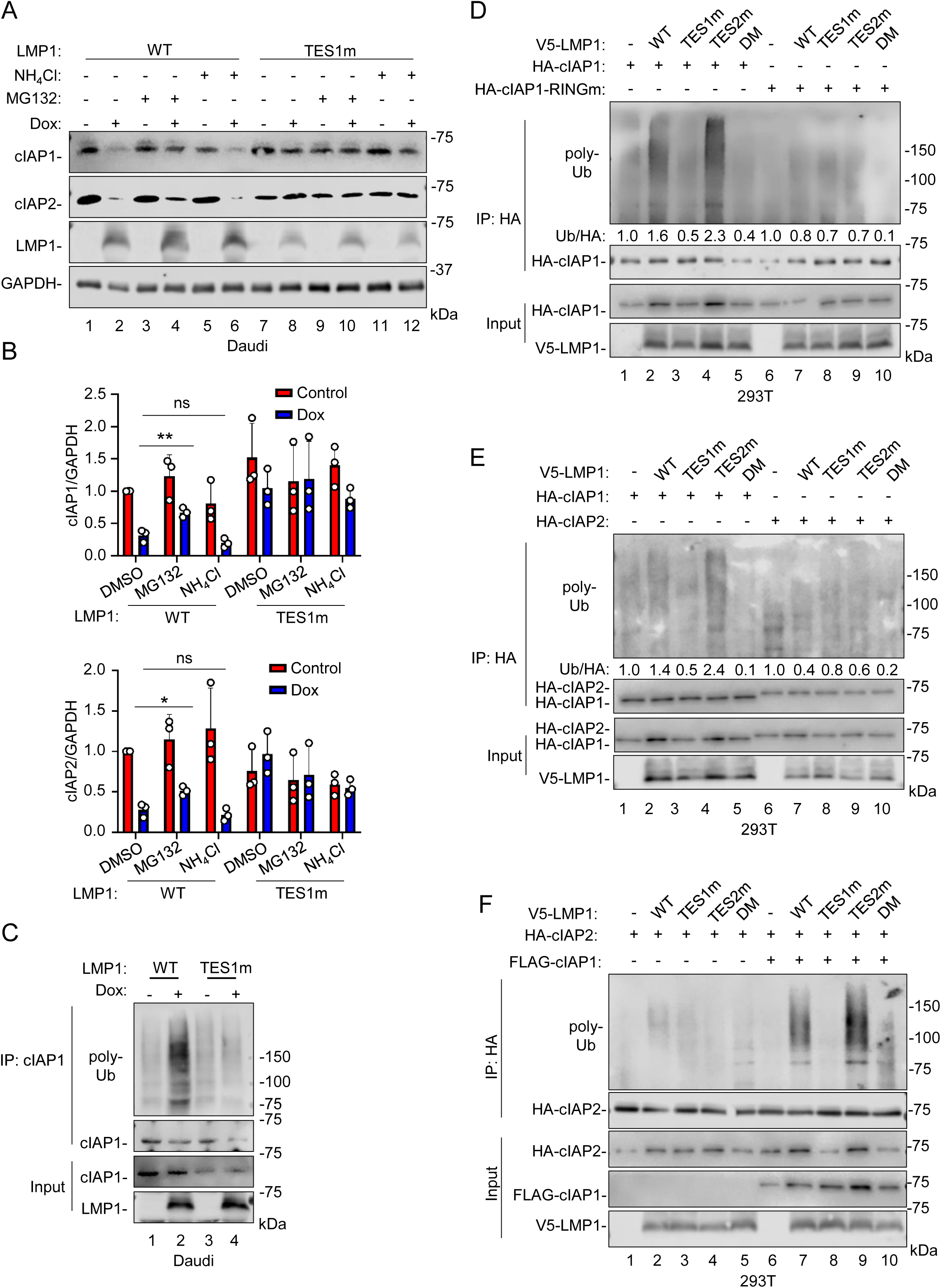
LMP1 triggers cIAP1/2 polyubiquitination and proteasomal degradation. (A) Analysis of proteasomal versus lysosomal roles in LMP1-driven cIAP turnover. Immunoblot analysis of WCL from Cas9+ Daudi cells induced for WT or TES1m LMP1 expression by 250 ng/mL Dox for 24 hours, followed by treatment with the proteasome inhibitor MG132 (5µM) or the lysosomal acidification inhibitor NH_4_Cl (20mM) for the last 8 hours. (B) Relative fold changes + SD of GAPDH-normalized cIAP1 or cIAP2 levels based on densitometry from n=3 replicates as in (A). Values in vehicle control treated cells uninduced for WT LMP1 were set to 1. Statistical significance was assessed by two-tailed unpaired Student’s t test (B). ns, not significant, *p<0.05, **p<0.01. (C) LMP1 TES1 signaling triggers cIAP1 polyubiquitination. Immunoblot analysis of 1% input vs endogenous cIAP1 immunopurified from Cas9+ Daudi cells induced for WT or TES1m LMP1 expression by 250 ng/mL Dox for 24 h. Cells were then treated with MG132 (5µM) for an additional 8 hours before collection. Cell lysates were boiled and immunoprecipitated with anti-cIAP1 antibody and protein A/G magnetic beads. (D) The cIAP1 RING domain is important for LMP1 TES1-driven cIAP1 polyubiquitination. Immunoblot analysis of input versus anti-HA-cIAP1, immunopurified from 293T cells that were transfected with expression vectors encoding HA-tagged cIAP1, cIAP1-RING mutant (RINGm, H588A) and empty vector or V5-tagged LMP1, as indicated. 24 hours following transfection, cells were treated with MG132 (5µM) for 24 hours prior to harvest. Cell lysates were boiled and immunoprecipitated with anti-HA-conjugated magnetic beads. Densitometry ratios of poly-ubiquitin versus HA-cIAP1 values are shown. (E) Analysis of LMP1-triggered cIAP1 vs cIAP2 polyubiquitination. Immunoblot analysis of input versus anti-HA-cIAP1 or anti-HA-cIAP2, immunopurified from 293T cells that were transfected with expression vectors encoding HA-tagged cIAP1 or cIAP2 and empty vector, V5-tagged LMP1, TES1m, TES2m or DM LMP1, as indicated. 24 hours following transfection, cells were treated with MG132 (5µM) for 24 hours prior to harvest. Cell lysates were boiled and then immunoprecipitated with anti-HA-conjugated magnetic beads. Ratios of polyubiquitin to HA-cIAP1 or cIAP2 IP are shown. (F) cIAP1 role in LMP1 TES1-driven cIAP2 poly-ubiquitination. Immunoblot analysis of input versus anti-HA-cIAP2, immunopurified from 293T cells that were transfected with expression vectors encoding empty vector, HA-tagged cIAP2, FLAG-tagged cIAP1, V5-tagged LMP1, TES1m, TES2m or DM LMP1, as indicated. 24 hours following transfection, cells were treated with MG132 (5µM) for 24 hours prior to harvest. Cell lysates were boiled and then immunoprecipitated with anti-HA-conjugated magnetic beads. Ratios of polyubiquitin to HA-cIAP2 IP are shown. Blots are representative of n=3 (A,D,E) or 2 (C and F) experiments. To achieve comparable cIAP1 protein levels given LMP1 boosting of the expression vector promoter activity, three times the amount of cIAP1 vector was transfected into cells lacking LMP1 expression or expressing the LMP1 TES1m or DM mutants (D-F).

cIAP1 and cIAP2 are E3 ubiquitin ligases, and their activity can play non-redundant versus redundant roles on downstream signaling pathways, in particular non-canonical NF-κB activation[82]. cIAP1 RING domain sequestration serves as an important mechanism of its E3 autoinhibition, which can be relieved by small molecule second mitochondria-derived activator of caspases (SMAC) mimetics[83-85]. We therefore next characterized how TES1 signaling alters cIAP1 vs cIAP2 poly-ubiquitination. Since the cIAP1 CARD-RING domains is able to downmodulate protein levels of both cIAP1 and cIAP2[86], we first examined whether LMP1 TES1 signaling triggers polyubiquitin chain attachment to cIAP1 molecules. To do so, we immunoprecipitated endogenous cIAP1 from Daudi cells mock-induced or induced for WT vs TES1m LMP1 for 24 hours, using whole cell lysates that were boiled for 5 minutes to disrupt protein-protein complexes. Intriguingly, abundant poly-ubiquitin chain modification was evident on cIAP1 immunoprecipitated from WT LMP1 expressing cells, but not from the mock induced or from TES1m expressing cells (**Fig. 5C**).

To build upon this result, we next co-expressed either HA-tagged wildtype or RING mutant cIAP1, in which an H588A point mutation abrogates cIAP1 activity[87]. 293T were then co-transfected with empty vector or expression constructs for WT, TES1m, TES2m or DM LMP1, together with HA-tagged wildtype vs RING mutant cIAP1 for 24 hours. Cells were then treated with MG132 for another 24 hours to inhibit proteasomal turnover. Anti-HA immunoprecipitation was performed on whole cell lysates that were boiled for 5 minutes, followed by immunoblot for poly-ubiquitin chains, which revealed that WT and TES2m LMP1, but not TES1m or DM LMP1, triggered abundant HA-cIAP1 poly-ubiquitin chain decoration. Interestingly, LMP1 failed to induce RING mutant HA-cIAP1 polyubiquitination (**Fig. 5D**).

We then cross-compared LMP1 TES1 and TES2 signaling effects on cIAP1 vs cIAP2 polyubiquitination. 293T were co-transfected with constructs encoding WT vs point mutant LMP1, together with HA-cIAP1 or cIAP2 expression vectors. MG132 was again added 24 hours post-transfection for a further 24 hours. HA-cIAP1 or cIAP2 were then immunoprecipitated from boiled whole cell lysates, and subject to polyubiquitin immunoblot. WT and TES2m LMP1 stimulated poly-ubiquitin chain attachment to HA-cIAP1, but to a considerably lesser degree to cIAP2, as judged by quantitation of the poly-ubiquitin to HA immunoblot signal (**Fig. 5E**). Therefore, our data suggests that TES1 signaling triggers cIAP1 polyubiquitination, potentially via autoubiquitination. In support of a key cIAP1 ubiquitin ligase role, co-expression of cIAP1 with cIAP2 triggered robust cIAP2 polyubiquitination in a TES1 dependent manner (**Fig. 5F**), indicating that cIAP1 likely supports LMP1-mediated polyubiquitination of cIAP2. While 293T also expresses endogenous cIAP1, we hypothesize that cIAP1 overexpression was required to stoichiometrically match transfected cIAP2 levels.

### TRAF3 is necessary for TES1-driven cIAP1 polyubiquitination

We hypothesized that TRAF3 is critical for cIAP1 polyubiquitination downstream of LMP1, given its role in cIAP1/2 recruitment to TES1. To test this, we first transfected 293T control versus TRAF2, 3 or 5 KO 293 cells with expression vectors encoding HA-cIAP1, alone or together with V5-tagged LMP1. We then immunoprecipitated HA-cIAP1 from boiled lysates, and by immunoblot observed robust cIAP1 polyubiquitination in lysates from LMP1 co-transfected control, TRAF2 KO or TRAF5 KO cells. By contrast, TRAF3 KO nearly completely ablated cIAP1 polyubiquitin chain decoration in LMP1-expressing cells (**Fig. 6A**). Similar results were observed in TRAF-edited Daudi Burkitt cells and LCLs (**Fig. 6B-C**). Taken together, these findings demonstrate that LMP1 induces proteasome-dependent degradation of cIAP1/2 through a mechanism that involves TES1-mediated recruitment of TRAF3 and cIAP1 ubiquitination.

**Figure 6.**
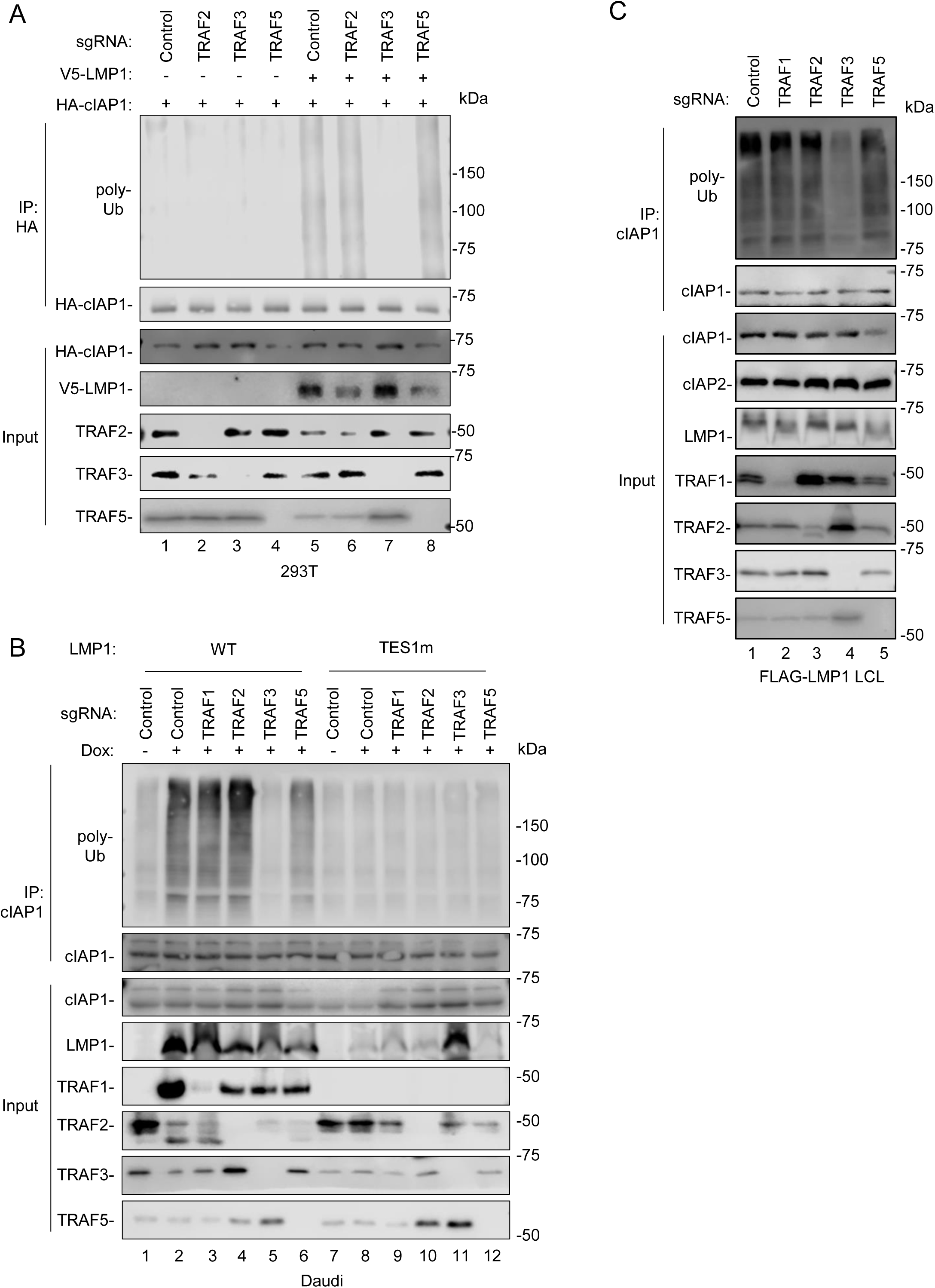
TRAF3 is required for LMP1-induced cIAP1 ubiquitination. (A) Analysis of TRAF roles in LMP-driven cIAP1 poly-ubiquitination in 293T cells. Immunoblot analysis of 1% input versus anti-HA-cIAP1, immunopurified from 293T cells that expressed control or TRAF targeting sgRNA and that were transfected with expression vectors encoding HA-cIAP1 and V5-LMP1 for 24 hours, as indicated. Cells were treated with MG132 5 µM) for 24 h prior to harvest. Three times the amount of cIAP1 vector was transfected into cells when not co-transfected with LMP1 to achieve comparable cIAP1 protein levels, to compensate for LMP1 boosting of the expression vector promoter activity. Cell lysates were boiled prior to immunoprecipitation with anti-HA magnetic beads. (B) Analysis of TRAF roles in LMP-driven cIAP1 poly-ubiquitination in Daudi B-cells. Immunoblot analysis of input versus cIAP1 immunopurified from Cas9+ Daudi cells that expressed control versus TRAF targeting sgRNAs and induced for LMP1 expression by 250ng/mL Dox for 24 hours. Cells were treated with 5µM MG132 for 24 hours prior to harvest. Lysates were boiled prior to immunoprecipitation with anti-cIAP1 antibody and protein A/G magnetic beads. (C) Analysis of TRAF roles in LMP-driven cIAP1 poly-ubiquitination in LCLs. Immunoblot analysis of input versus cIAP1 immunopurified from Cas9+ Daudi cells that expressed control versus TRAF targeting sgRNAs. Cells were treated with 5µM of MG132 for 24 hours prior to harvest. Cell lysates were boiled and immunoprecipitated with anti-cIAP1 antibody and protein A/G magnetic beads. Anti-mouse IgG was used as a negative control. Blots are representative of n = 2 experiments.

### cIAP1 or cIAP2 over-expression impair LMP1 mediated non-canonical NF-κB activity

Non-canonical NF-κB pathways culminate in processing of the p100 NF-κB precursor into the active p52 transcription factor subunit. The observation that LMP1 induces degradation of cIAP1/2 led us to hypothesize that cIAP1 loss is essential for LMP1-mediated activation of non-canonical NF-κB signaling, akin to how SMAC mimetics activate non-canonical NF-κB[84]. To test the model that cIAP1 loss is important for TES1 mediated non-canonical activity, we stably expressed control GFP versus cIAP1 in Daudi cells, and then mock induced or induced LMP1 expression for 24 hours. Immunoblot analysis of WCL revealed that enforced cIAP1 expression significantly diminished LMP1 driven p100:p52 processing (**Fig. 7A-B**). Likewise, LMP1-induced p100:p52 processing was significantly impaired by co-expression of either cIAP1 or cIAP2 in 293T cells (**Fig. 7C-D**), consistent with a model in which TES1 driven turnover of both cIAP1 and cIAP2 serves as a major driver of downstream non-canonical NF-κB activity. Collectively, our data is consistent with a model in which LMP1 targets TRAF2, cIAP1 and cIAP2 for proteasomal degradation, which results in activation of downstream non-canonical NF-κB activity and release of active p52 transcription factor subunits for modulation of nuclear target gene expression (**Fig. 8**).

**Figure 7.**
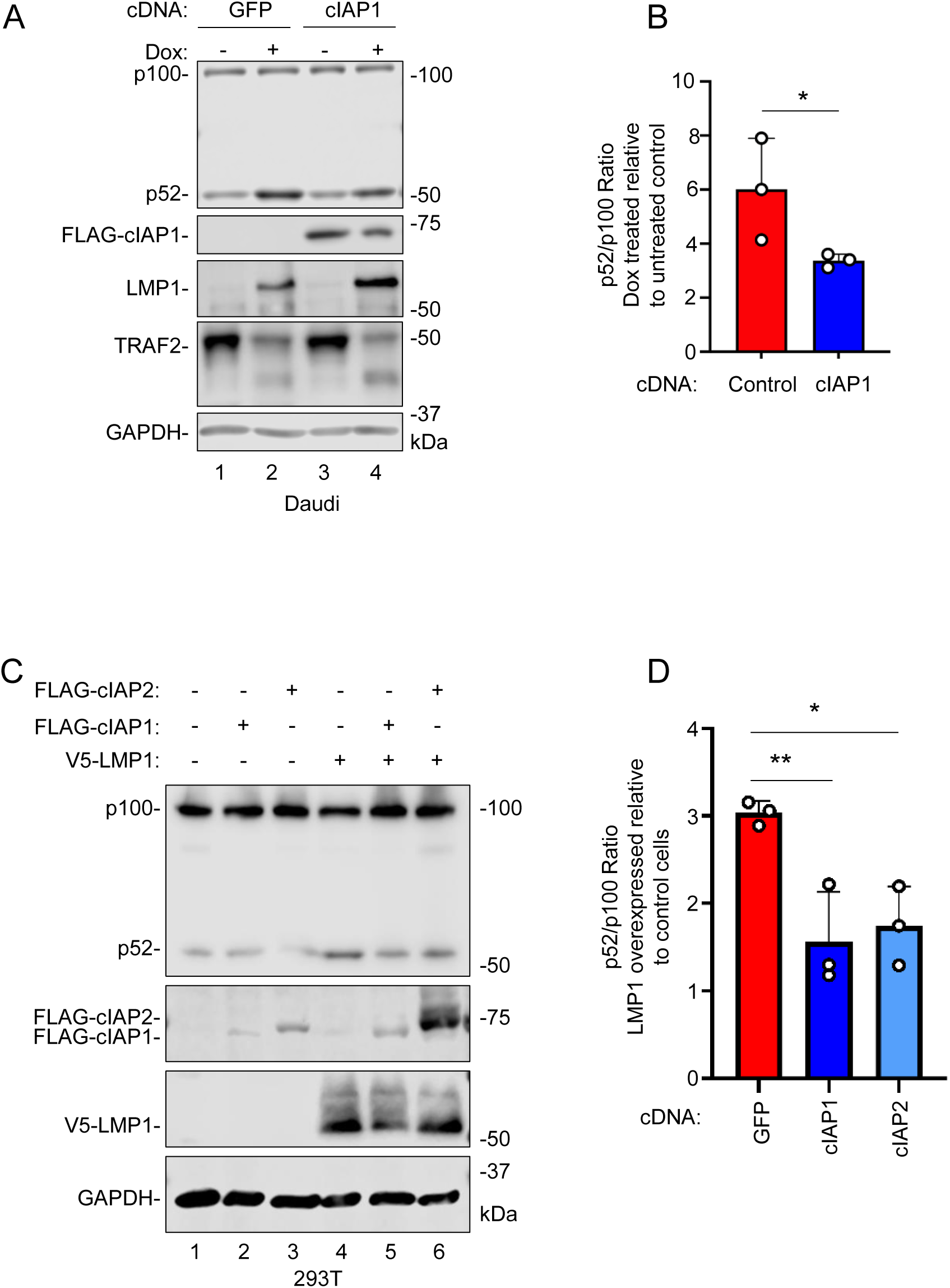
cIAP1/2 overexpression impairs LMP1-induced non-canonical NF-κB pathway activation. (A) Effect of cIAP1 over-expression on LMP1-induced p100 processing. Immunoblot analysis of WCL from Daudi cells transfected with control GFP versus cIAP1 expression vectors for 24 hours and then mock induced or induced for LMP1 expression with 250 ng/mL Dox for 24 hours, as indicated. (B) Quantification of p100:52 processing from n=3 immunoblots as in (A). Shown are relative p52:p100 ratios + SD, indicative of non-canonical NF-κB activity, from n=3 replicates. (C) cIAP1 or cIAP2 overexpression effects on LMP1-induced non-canonical activity. Immunoblot analysis of WCL from 293T cells transfected with expression vectors encoding empty vector control, FLAG-cIAP1, FLAG-cIAP2 or V5-LMP1, as indicated for 24 hours. (D) Quantification of p100:p52 processing from n=3 immunoblots as in (C). Shown are relative relative p52:p100 ratios + SD from n=3 replicates from cells transfected with LMP1 vs with empty vector control for each condition (i.e. lane 2 vs 1, lane 4 vs 3, etc). Statistical significance was assessed by two-tailed unpaired Student’s t-test (B) or one-way ANOVA followed by Tukey’s multiple comparisons test (D). ns, not significant, *p<0.05, **p<0.01. Blots are representative of n=3 experiments.

**Figure 8.**
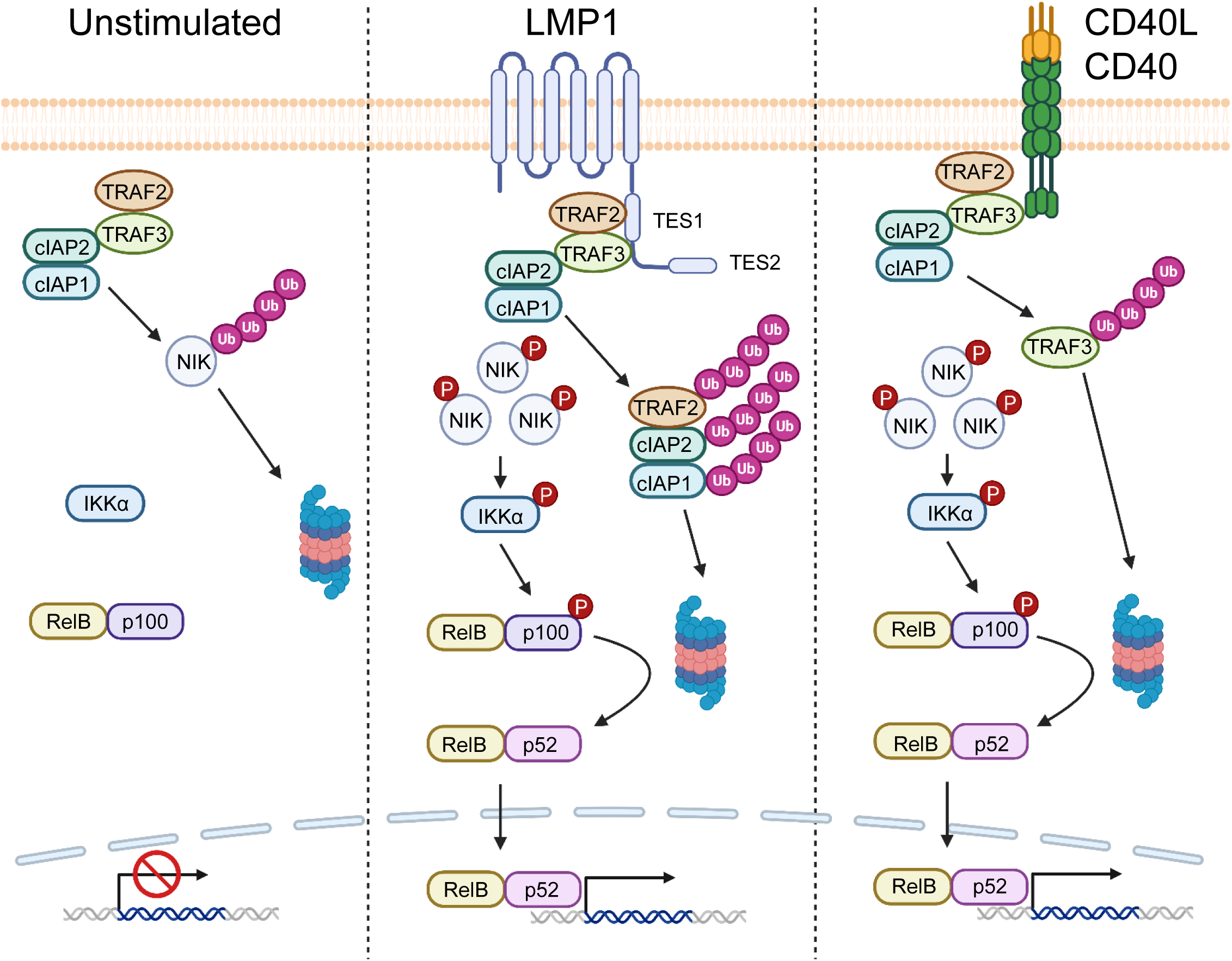
Schematic model of LMP1 versus CD40 driven non-canonical pathways. Under unstimulated conditions, the TRAF-cIAP complex targets the kinase NIK for proteasomal degradation, thereby preventing non-canonical NF-κB signaling. LMP1 TES1 triggers TRAF2, cIAP1 and cIAP2 for degradation, which initiates downstream signaling and p100:p52 processing. By comparison, CD40L/CD40 signaling induces degradation of TRAF3 to drive downstream non-canonical NF-kB pathway activation.

## Discussion

Immune receptors and viral proteins utilize multiple distinct mechanisms to initiate non-canonical NF-κB signaling, which is critical for B-cell differentiation and survival. Here, we present multiple lines of evidence that LMP1 TES1 signaling, which is critical for EBV-mediated primary B cell immortalization and for lymphoblastoid B-cell survival[25-29, 62], triggered rapid depletion of the E3 ubiquitin ligases cIAP1, cIAP2 and TRAF2. This phenomenon was dependent on TRAF3, which binds tightly to a TES1 PxQxT motif[25-29, 50, 53, 75], and which we now report in turn recruits cIAP1 and cIAP2 to LMP1. LMP1 and TRAF3 together activated cIAP1 RING dependent ubiquitination and degradation of cIAP1 and cIAP2 as well as of TRAF2. Loss of cIAP1/2 drove downstream p100 processing to p52, whereas enforced cIAP1 expression impaired LMP1-driven non-canonical activation.

It has been proposed that LMP1 activates non-canonical NF-κB signaling pathway by sequestering TRAF3, thereby preventing TRAF3-mediated degradation of NIK[53]. However, quantitative data suggests that only ∼50% of TRAF3 is associated with LMP1 in LCLs[48, 50, 51], raising the question of whether sequestration is sufficient to robustly activate the pathway. In addition to tightly binding to TRAF3, our findings complement this prior study by now suggesting that LMP1 evolved a mechanism related to, but distinct from that utilized by CD40 and BAFF receptors to initiate non-canonical activity. Rather than targeting cIAPs directly, CD40 and BAFF receptors target TRAFs 2 and 3 for proteasomal degradation[65, 66], which serves to interrupt cIAP repression of the kinase NIK to induce p100:p52 processing. LMP1 instead utilized TRAF3 to target cIAP1, cIAP2 and TRAF2 for degradation.

TRAF3 was originally identified as a NIK-binding protein in a yeast two-hybrid screen[88]. In the absence of stimuli, TRAF3 serves as an adaptor protein that enables cIAP1/2 to target NIK for ubiquitination and proteasomal degradation to prevent non-canonical NF-κB activation[43]. NIK ubiquitination involves a multi-subunit ubiquitin ligase complex composed of TRAF2, TRAF3, cIAP1 and/or cIAP2. Within this complex, TRAF2, but not TRAF3, directly interacts with cIAP1/2[71]. Since cIAPs, as part of the TRAF–cIAP complex, are responsible for ubiquitinating and degrading NIK under resting conditions, their elimination disrupts this suppressive complex to activate non-canonical NF-κB signaling. Thus, in addition to TRAF3 sequestration, degradation of cIAP1/2 represents a critical step that amplifies LMP1-mediated activation of the noncanonical NF-κB pathway.

TRAFs 1, 2, 3 and 5 each associate with LMP1 TES1 within TRAF homo- or heterotrimeric complexes. Of these, TRAF3 binds most tightly to LMP1[48, 50, 51, 53]. In LCLs, only ∼5% of the TRAF2 pool is associated with LMP1, in contrast with the ∼50% of TRAF3 that is bound by LMP1[48, 50, 51]. Furthermore, even though TRAF2 binds directly to cIAP1/2[71, 89], our data suggest that TRAF2 alone is not sufficient to recruit cIAPs to LMP1, since LMP1 targeted both cIAP1 and cIAP2 for degradation in TRAF2 knockout cells. It remains plausible that TRAF3-bound LMP1 serves as a scaffold for TRAF2/cIAP1/cIAP2 complexes, which are then routed to the proteasome. In support, we found that TRAF3 KO abrogated cIAP1/2 binding to LMP1. Alternatively, it is possible that functional redundancy between TRAFs 2 and other TRAFs, in particular TRAF5, may enable cIAP1/2 recruitment to LMP1 in a manner that is not absolutely dependent on TRAF2.

A future goal will therefore be to identify how LMP1-bound TRAF3 activates cIAP1 ubiquitin ligase activity, autoubiquitination and proteasomal degradation. Small molecules that promote cIAP degradation called SMAC mimetics activate the non-canonical NF-κB pathway[90]. In some respects, TES1 signaling is reminiscent of SMAC mimetics, whose binding to cIAPs triggers rapid autoubiquitination and depletion. As observed with SMAC mimetics, cIAP depletion is dependent on cIAP1 RING ubiquitin ligase activity[91]. However, in contrast to SMAC, which similarly activates both cIAP1 and 2, our data suggest that cIAP1 ligase activity is critical and targets both cIAP1 and cIAP2 for degradation.

cIAP1 and cIAP2 are themselves NF-κB pathway targets and their mRNA abundances are upregulated by LMP1[29, 92, 93]. In latency III B cells such as LCLs, LMP1-driven NF-κB activity drives cIAP1 and 2 re-syntheses. Thus, cIAP1/2 are continually expressed and degraded in LMP1+ cells. Consequently, they can be detected at steady state levels by immunoblot of cells with stable LMP1 expression. However, CHX chase analysis revealed shorter T_1/2_ in the presence of TES1 signaling. This may serve to sufficiently deplete local pools of cIAPs in order to remove their brake on the non-canonical NF-κB pathway, while leaving some residual pools to perform other essential cellular roles, such as in support of pro-survival signaling downstream of the TNF receptor[92, 94, 95]. In support of this point, LMP1 signals constitutively and independently of ligand, in contrast to CD40, and EBV may have therefore needed to evolve a mechanism to downmodulate without completely suppressing cIAP expression in cells that express LMP1 for long durations.

It is difficult to prove that TRAF2 is dispensable for TES1 signaling, since its depletion induces non-canonical NF-κB activity. Interestingly however, we observed the appearance of a lower-molecular-weight band immunoreactive with anti-TRAF2 antibody upon conditional LMP1 expression, present also in LCLs and in MUTU III. This may represent a TRAF2 isoform or more likely a TRAF2 degradation product. We previously reported that LMP1 triggers K63-linked polyubiquitin chain attachment to TRAF2[28], but this is typically associated with signal transduction rather than degradation. LMP1 may therefore induce partial TRAF2 cleavage in a manner not dependent on proteasome activity. A future objective will be to identify the protease that LMP1 directs to TRAF2, as well as the effect of this cleavage event on other TRAF2 activities.

Our data reveal that LMP1 promotes the ubiquitination and degradation of cIAP1 via a mechanism dependent on cIAP1’s intrinsic E3 ligase activity. This observation suggests that LMP1 may function similarly to small-molecule antagonists such as SMAC mimetics, which trigger cIAP autoubiquitination and degradation[83-85]. However, an important future object will be to define how LMP1 TES1 signaling activates cIAP1, but not cIAP2 polyubiquitination. Understanding how LMP1 specifically engages cIAP1’s E3 ligase activity may reveal new details about IAP regulation and provide potential therapeutic targets in EBV-associated disorders. In particular, further molecular insights may allow the development of rational small molecules that perturb either LMP1 recruitment of TRAF3 and/or LMP1/TRAF3-mediated cIAP1/2 degradation. These would have the potential to selectively block LMP1, without inhibiting immunoreceptor signaling, including by CD40.

In conclusion, we identified that LMP1 TES1 activates the non-canonical NF-κB pathway through targeted degradation of cIAP1/2 and TRAF2. This mechanism involves the assembly of a multi-protein complex mediated by LMP1 TES1 and TRAF3 and drives downstream p100 processing to the active p52 NF-κB transcription factor subunit. These findings suggest that it may be possible to develop small molecule inhibitors that selectively block the apparently unique mechanism by which LMP1 TES1 activates non-canonical NF-κB activity, which is critical for EBV-mediated B cell transformation into immortalized lymphoblastoid cells and also for their survival.

## Acknowledgements

This work was supported by NIH R01 CA228700 to BEG. YS was supported by American Cancer Society Post-doctoral Fellowship PF-23-1144614-01-IBCD. BM was supported by American Cancer Society Post-doctoral Fellowship PF-24-1250090-01-IBCD. SL was supported by the Harvard University Center for AIDS Research (CFAR) NIH funded program (P30 AI060354), which is supported by the following NIH Co-Funding and Participating Institutes and Centers: NIAID, NCI, NICHD, NHLBI, NIDA, NIMH, NIA, NIDDK, NIMHD, NIDCR, NINR, OAR, and FIC. We thank Makoto Ohashi and Eric Johannsen for valuable suggestions on the EBV BAC system. We thank Greg Smith for sharing the BM2710 bacteria and for valuable suggestions.

## Material and Methods

### Cell lines

HEK-293T were purchased from ATCC and cultured in Dulbecco’s modified Eagle medium (DMEM, Gibco) supplemented with 10% fetal bovine serum (FBS, Gibco) and 1% penicillin/streptomycin (Gibco). Daudi and Akata Burkitt cells were obtained from Elliott Kieff. MUTU I and MUTU III Burkitt cells were obtained from Jeff Sample. FLAG-LMP1 LCLs were obtained from Elliott Kieff, generated using a recombinant EBV in which the FLAG tag encoding DNA sequence was knocked immediately upstream of the LMP1 ATG initiator codon[50]. All B cell lines were cultured in Roswell Park Memorial Institute (RPMI) 1640 (Life Technologies), supplemented with 10% v/v FBS and 1% pen/strep. B cell lines with stable *Streptococcus pyogenes* Cas9 expression as previously described[98] were maintained with 5 μg/mL blasticidin (InvivoGen). Daudi and Akata cell lines with doxycycline-inducible WT, TES1m, TES2m, or DM LMP1 alleles were previously described[29]. Conditional LMP1 cells were maintained in either 50 μg /mL hygromycin or 3 μg/mL puromycin. For LMP1 induction studies, cells were seeded at 0.5 × 10⁶ cells/mL and treated with 250 ng/mL doxycycline (Sigma) for 24 hours. All cells were incubated at 37°C with 5% CO_2_.

### Primary Human B cells

Deidentified and discarded leukocytes, left over following voluntary platelet donation, were obtained from the Brigham and Women’s Hospital Blood Bank following receipt of informed consent by the blood bank. Our non-human subjects research studies with these discarded cells were done under approved Institutional Review Board protocols. Peripheral blood mononuclear cells (PBMCs) were isolated using Lymphoprep Density Gradient Medium (Stem Cell Technologies), followed by negative selection of primary B cells using RosetteSep and EasySep Human B Cell Enrichment Kits (Stem Cell Technologies), in accordance with the manufacturers’ instructions. Primary B-cells were grown in RPMI-1640 supplemented with 10% FBS.

### Construction of bacterial artificial chromosome (BAC) EBV LMP1 mutant genomes

The EBV p2089 BACmid was used for production of recombinant EBV genomes with wildtype, TES1 mutant or TES2 mutant genomes, as previously described[67]. P2089 encodes the EBV B95.8 strain genome, as well as a cassettes encoding the prokaryotic F-factor, kanamycin resistance cassette, green fluorescence protein and Hygromycin B resistance cassette, as described previously[99]. BACmids were constructed using the GS1783 E. coli–based En Passant method previously described[100]. TES1 and TES2 point mutations were constructed using established BAC engineering protocols. For the TES1m EBV, 204 PQQAT was mutated to AQAAA to inactivate TES1 signaling. For the TES2m EBV, 384YYD was mutated to ID to inactivate TES2 signaling. BAC genome fidelity was validated by EcoRI, BamHI and NcoI DNA restriction map analysis, by DNA Sanger sequencing of high fidelity PCR-amplified TES regions, and by demonstration that revertants were fully transforming. BACmids were isolated and electroporated into spectinomycin-resistant BM2710 E. coli using 0.1cm cuvette at 1.5kV, 200 Ohms, 25µF for infection of 293 cells. 293 producer cells were established by BM2710 *E. coli* invasion, as previously described[101]. Single cell 293 producer clones were screened.

### EBV stock preparation and quantification

EBV-positive 293 producer cells were reverse-transfected with expression plasmids encoding BZLF1 and BALF4 to induce the lytic cascade, as previously described[67]. Three days post transfection, supernatants were harvested and centrifuged at 1200 rpm for 10 mins and then at 4000 rpm for 10 mins to remove cell debris. Supernatants were passed through a 0.45µm filter to remove debris and concentrated by ultracentrifugation at 25000 rpm for 2 hours at 4°C. Viral stocks were tittered by the green Raji Assay[102]. In brief, 1 × 10⁵ Raji cells were seeded in 100 μL per well of a 96-well plate and infected with serial dilutions of EBV. The percentage of GFP⁺ Raji cells was determined by flow cytometry 2 days post-infection. The concentration of infectious EBV virions was calculated as Green Raji Units per mL. For infection experiments, EBV was added at a multiplicity of infection (MOI) of 0.1 GRU per target cell.

### Antibodies and Reagents

For immunoblot analysis, Cell Signaling Technology antibodies against cIAP1 (D5G9), cIAP2 (58C7), XIAP (3B6), TRAF1 (45D3), TRAF3 (#4729), TRAF5 (#D3E2R), FLAG (M2, #2368), V5 (#D3H8Q), NIK (#4994) and ubiquitin (P4D1, #3936) were used at 1:1000 for immunoblot analysis. Antibodies against TRAF2 (Proteintech 26846-1-AP), TRAF2 (Invitrogen SD205-06), HA (Abcam ab9110), and p100/52 (EMD Millipore #05-361) were used at 1:1000. Anti-GAPDH (EMD Millipore #MAB374) was used at 1:5000. The anti-LMP1 mouse monoclonal antibody S12[103] was used from hybridoma supernatant at 1:1000. Cell Signaling Technology HRP-linked anti-mouse (#7076) and anti-rabbit (#7074) antibodies were used as secondary antibodies for immunoblot analysis at 1:5000. Proteintech anti-cIAP1 (1H3F1) and Abcam anti-HA (ab9110) were used for immunofluorescence assay at 1:200. Anti-FLAG M2 (SIgma) and Proteintech anti-cIAP1 (1H3F1) were used for immunoprecipitation at 1:200. 100 µg/mL Cycloheximide (R&D systems), 5 µM MG132 (SelleckChem), 20 mM ammonium chloride (sigma) and 100 ng/ml CD40L (Enzo Life Sciences) were used for cell stimulation.

### CRISPR-Cas9 editing

CRISPR/Cas9 editing in B-cell lines with stable Cas9 expression was performed as previously described[62, 98]. Briefly, Broad Institute pXPR-510 control sgRNA (targets a non-coding intergenic region), Avana, or Brunello sgRNA, as listed in **Table 1**, were cloned into pLentiGuide-puro (a gift from Feng Zhang, Addgene plasmid #52963), pLenti-spBsmBI-sgRNA-Hygro (a gift from Rene Maehr, Addgene plasmid #62205), or pLentiGuide-zeo (a gift from Rizwan Haq, Addgene plasmid #160091). The presence of sgRNA sequences were confirmed by Sanger sequencing. To generate lentivirus containing the sgRNA, 293T cells were transfected with pCMV-VSV-G (a gift from Bob Weinberg, Addgene plasmid #8454), psPAX2 (a gift from Didier Trono, Addgene plasmid #12260), and the sgRNA expression vector using the TransIT-LT1 transfection reagent (Mirus Bio). 293T supernatants were harvested and added to target B-cells at 48 and 72 hours post-293T transfection. As a control, target cells were transduced with sgRNA against GFP (pXPR-011, a gift from John Doench). Transduced cells were selected with puromycin (3 μg/ml), zeocin (200 μg/mL) or hygromycin (300 μg/mL). On-target CRISPR effects were validated by immunoblotting.

**Table 1.**
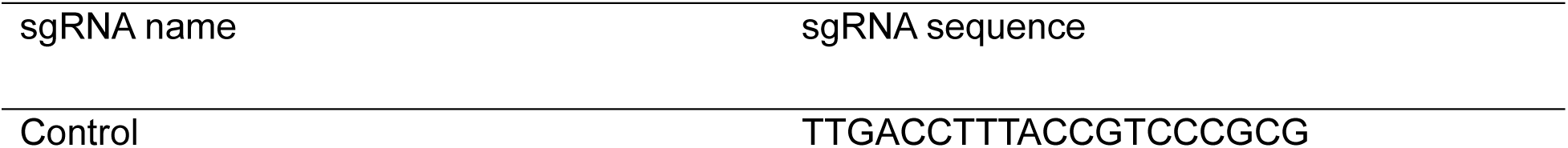
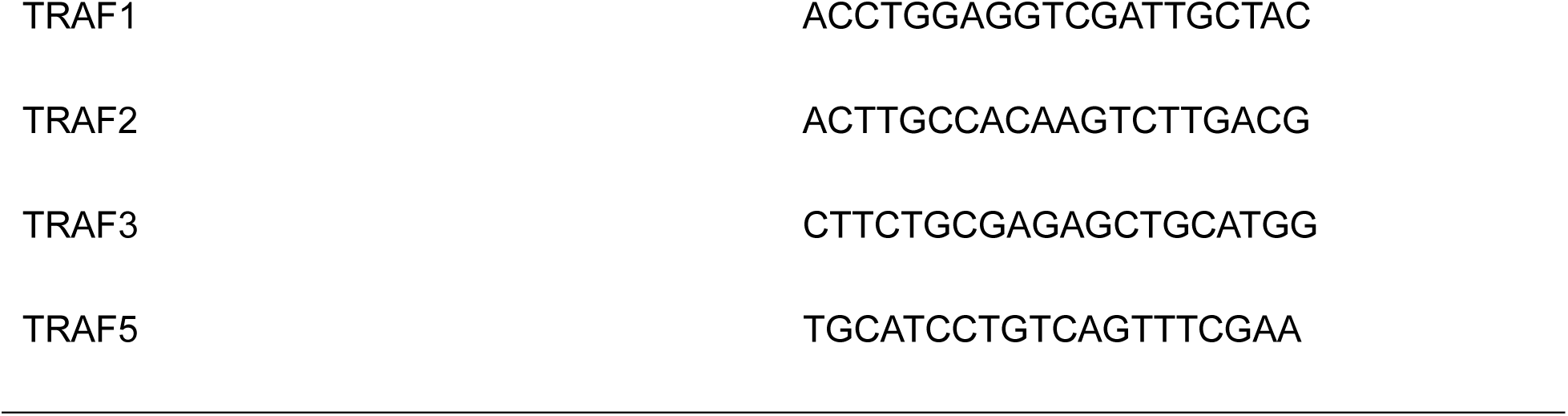
sgRNA sequence used in this study.

For CRISPR/Cas9 editing in HEK-293T cells, Brunello sgRNAs were cloned into pSpCas9(BB)-2A-Puro (PX459) V2.0 (a gift from Feng Zhang Addgene plasmid # 62988), which containing both Cas9 and cloning sites for sgRNA. Successful sgRNA ligation was confirmed by Sanger sequencing. 293T cells were transfected with the sgRNA-expressing PX459 vector using the TransIT-LT1 transfection reagent. Cells were not subjected to puromycin selection. Forty-eight hours post-transfection, cells were collected and on-target CRISPR effects were assessed by immunoblotting.

### SDS-PAGE and immunoblot

Immunoblot analyses were performed as described previously[104]. In brief, whole cell lysates were resolved by SDS-PAGE and transferred to nitrocellulose membranes (Bio-Rad), blocked with 5% nonfat dry milk in TBST for 1 h and incubated with primary antibodies at 4 °C overnight. Blots were washed three times in TBST and incubated with secondary antibodies for 1 h at room temperature. Blots were washed three times in TBST, incubated with ECL (Thermo Fisher #34578) and imaged by the Licor Fc platform. For quantification, densitometry data was obtained using the Image Studio Lite Ver 5.2 program.

### Immunoprecipitation

Immunoprecipitation was performed as previously described[105]. For B cells, 80 – 150 million cells were lysed in 1% v/v NP40, 150 mM Tris, 300 mM NaCl, 1 X cOmpleteTM EDTA-free protease inhibitor cocktail (Sigma), 1 mM Na3VO4 (Sigma) and 1 mM NaF (Sigma). 10% (v/v) of an aliquot from each lysate was preserved as the input. The rest of the lysate was incubated with a specific antibody targeting the epitope of interest overnight at 4°C with gentle rotation, followed by incubating with Protein A/G magnetic beads for 2 hours. For anti-HA tag IP, 20 uL magnetic HA beads (Thermo Fisher) were added to the whole cell lysate. Following multiple washes with lysis buffer, proteins were eluted by boiling at 95 °C for 10 mins.

### Ubiquitination analysis

Cells were pretreated with MG132 for 24 hours before collection. 150 million B cells or 100 million HEK-293T cells were used. For HEK-293T cells, cells were transduced to express LMP1, cIAP1, and cIAP2 prior to MG132 treatment. To achieve similar levels of exogenous cIAP1/2 expression, one-third the amount of cIAP1/2 vectors was used in co-transductions with LMP1 compared to transductions performed without LMP1 or with LMP1 TES1m or DM. Site-directed mutagenesis using the NEB Q5 Site-Directed Mutagenesis Kit (E0554) was used to make the RING domain mutation in cIAP1. Cells were lysed with lysis buffer containing 1% v/v NP40, 150 mM Tris, 300 mM NaCl, 1 X cOmpleteTM EDTA-free protease inhibitor cocktail (Sigma), 1 mM Na_3_VO_4_, 1 mM PMSF (Thermo Fisher), 4 mM 1,10 o-phenanthroline (Sigma), 2 mM sodium pyrophosphate (Sigma) and 1 mM EDTA (Life Technology). 10% (v/v) of an aliquot from each lysate was preserved as the input. The remaining cell lysates were boiled at 95 °C for 5 mins, and then incubated with the specific antibody targeting the protein of interest overnight at 4°C with gentle rotation, followed by incubating with Protein A/G magnetic beads for 2 hours. Beads were washed with cold lysis buffer for four times, and then eluted by incubating at 95 °C for 10 mins. The eluted proteins were then subjected to immunoblot analysis.

### Inhibition of Protein Degradation

To inhibit proteasome-mediated protein degradation in LMP1 conditional expression cells, cells were treated with 5 μM MG132 for 8 hours following 24 hours of LMP1 induction. For inhibition of lysosome-mediated protein degradation, 20 mM ammonium chloride was added under the same conditions as MG132 treatment.

### Immunofluorescence analysis

Cells dried on glass slides were fixed in 4% paraformaldehyde (Santa Cruz) in PBS for 10 minutes, followed by permeabilization with 0.5% Triton X-100 (Sigma) in PBS for 5 minutes. Cells were blocked with 1% bovine serum albumin (Sigma) in PBS for 1 hour at room temperature. Cells were incubated with primary antibodies (1:200 dilution) in blocking buffer overnight at 4 °C. Following two washes with PBS, cells were stained with secondary antibodies. For detecting cIAP1 and LMP1, a cocktail of secondary antibodies conjugated with fluorophores were used with 1:1000 dilution in DPBS. For cIAP1, Goat anti-Mouse IgG Alexa Fluor™ 568 (H+L) secondary antibody (Invitrogen, # A-11031) was used, and for LMP1-HA, Donkey anti-Rabbit IgG Alexa Fluor™ 647 (H+L) secondary antibody (Invitrogen, # A-31573) was used. Cells were then washed twice with PBS and mounted overnight in ProLong™ Gold Antifade Mountant with DAPI (Thermo Fisher). Imaging and analysis were carried out using a Zeiss LSM 800 confocal microscope and Zeiss Zen Lite (Blue) software, respectively. To quantify the number of cells in which LMP1 and cIAP1 signals were colocalized, the following methods were used. The total number of cells in each field of view was first determined by single channel automated counting of the DAPI stained nuclei. Each field typically contained 10-20 cells. Colocalization analysis is performed by plotting the fluorescence intensity of each channel as a function of distance along the line. Colocalization of LMP1 and cIAP1 was determined by comparing the fluorescence intensity curves where overlapping of two peaks was judged as colocalization. Colocalization was then confirmed by ImageJ with the Colo2 plug-in. The % cells which have LMP1/cAIP1 colocalization was calculated by # cells which have colocalization / # total cells in the field.

### Statistical analysis

All experiments were performed with two or three independent experiments. Statistical significance was assessed with Student’s t test using GraphPad Prism 9 software, where NS = not significant, p > 0.05; * p < 0.05; ** p < 0.01; *** p < 0.001.

### Schematic Models

BioRender was used to create all schematic models.

---

**Fig. S1.**
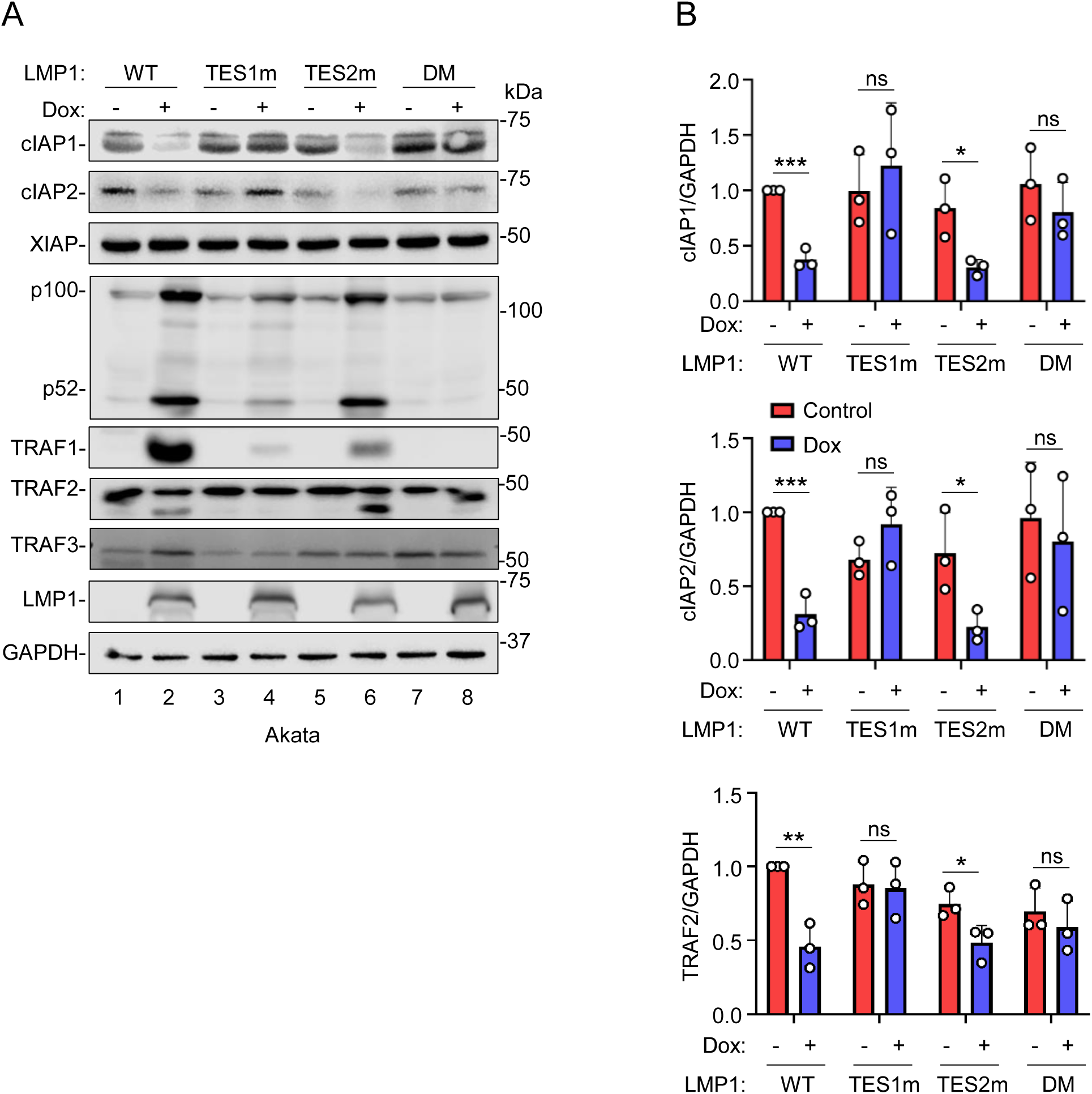

**Fig. S2.**
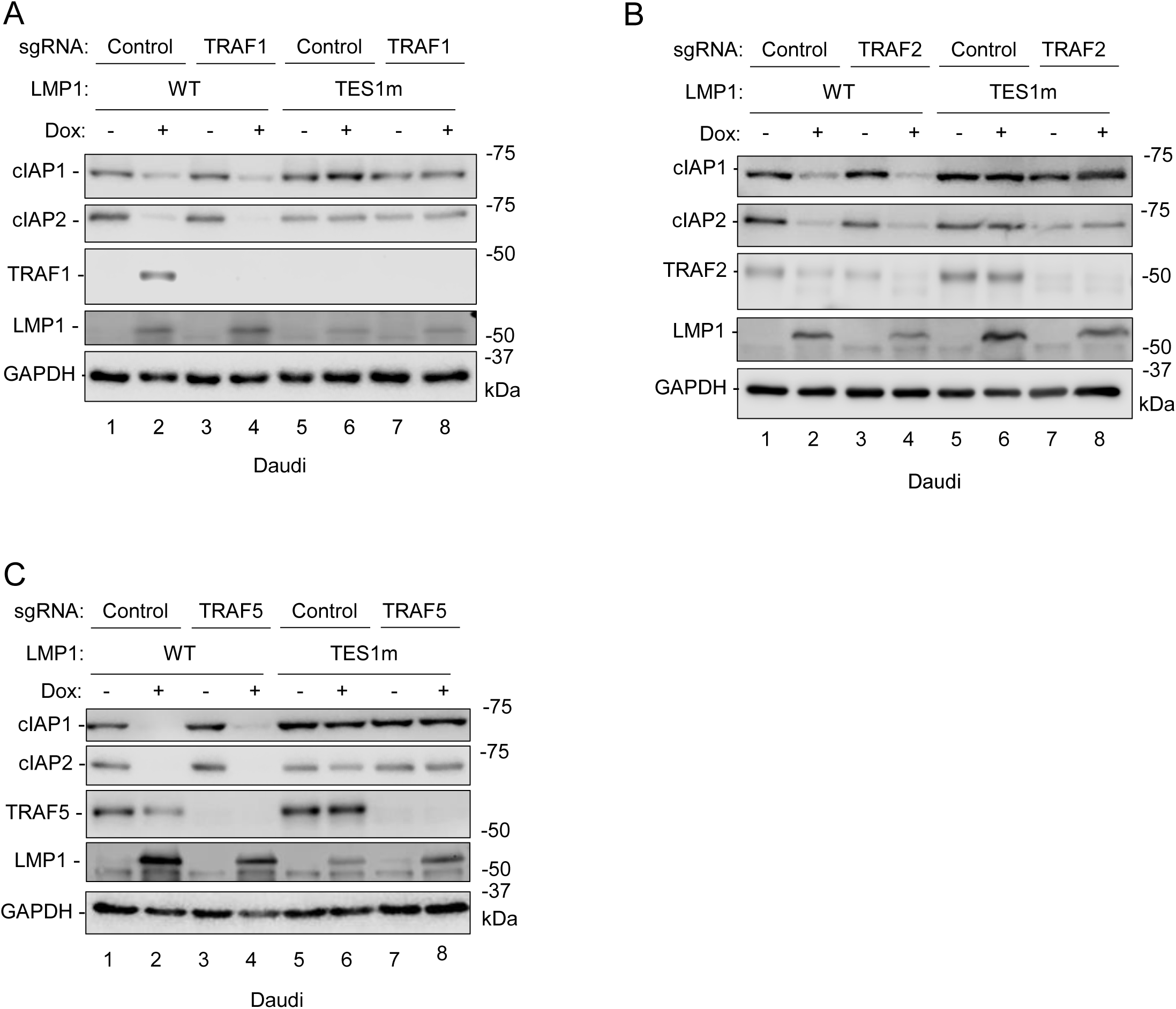

**Fig. S3.**
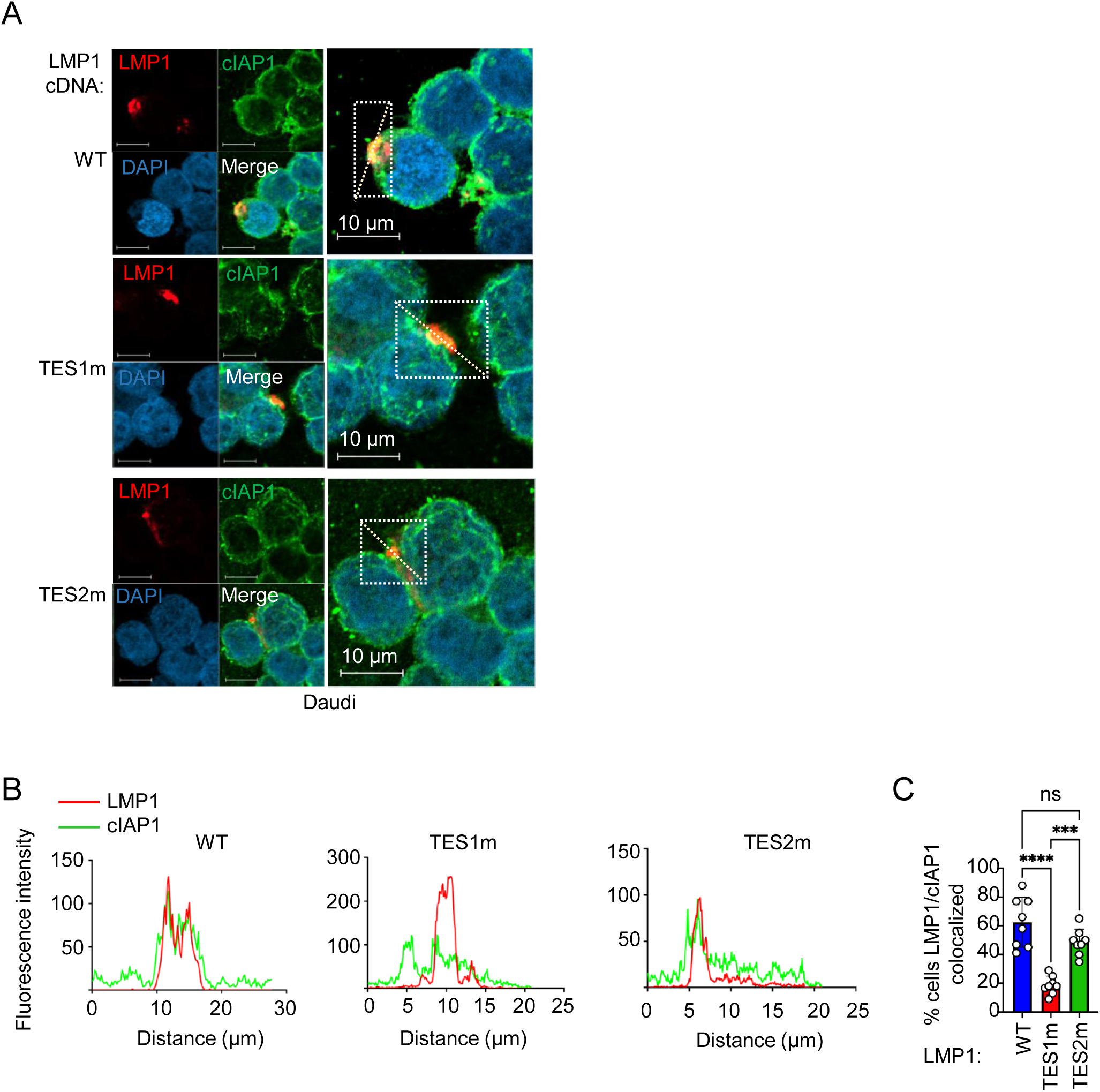

**Fig. S4.**
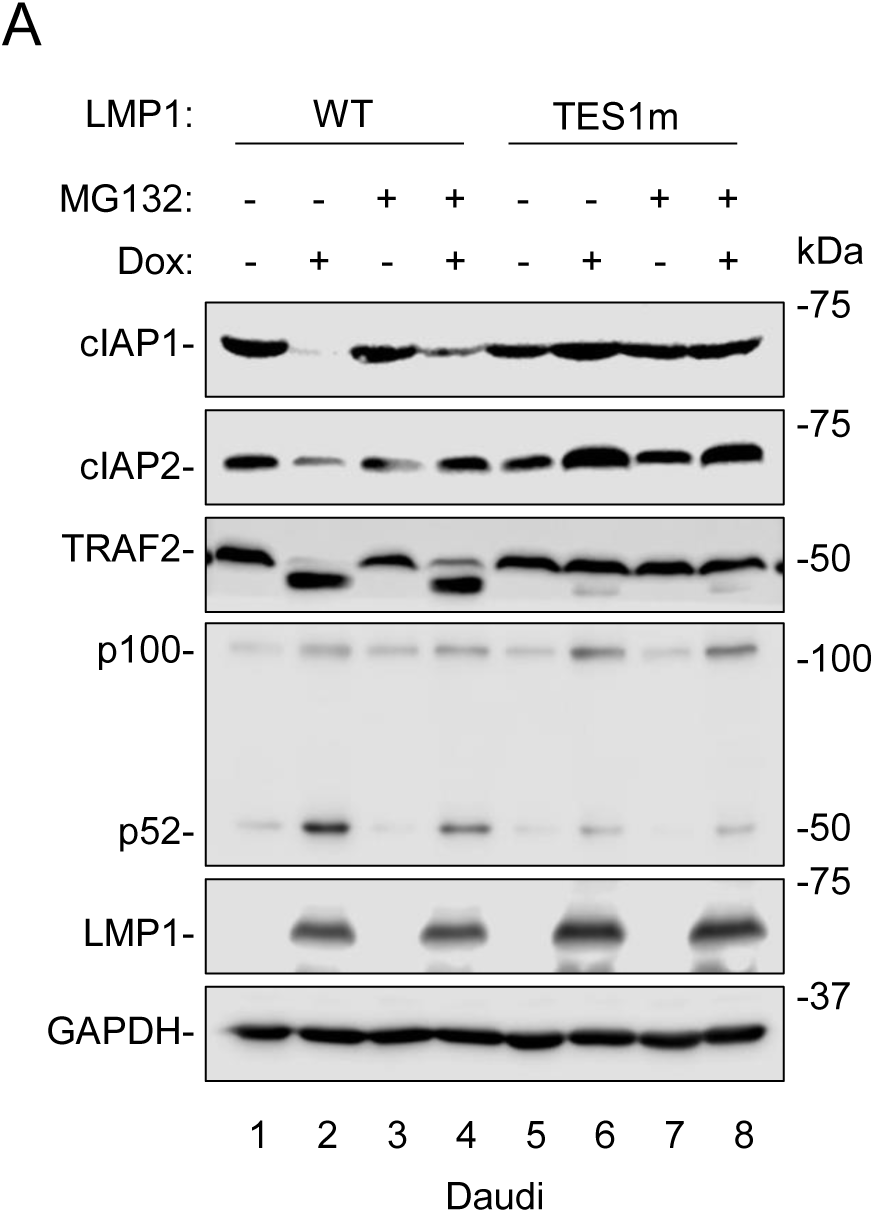

## Notes

### Competing Interest Statement

The authors have declared no competing interest.

