## Supplemental materials for "Epstein-Barr Virus Latent Membrane Protein 1 targets cIAP1, cIAP2 and TRAF2 for Proteasomal Degradation to Activate the Non-canonical NF-κB Pathway"

**Short title: LMP1 Targets cIAP1/2 and TRAF2 for Non-canonical NF-κB Activation**

Yizhe Sun[1,2*#], Shunji Li[1,2#], Bidisha Mitra[1,2], Ling Zhong[1,2], Aretina Zhang[1,2] and Benjamin E. Gewurz[1-2*]

[1]Division of Infectious Diseases, Department of Medicine, Brigham and Women's Hospital, Boston, Massachusetts, United States of America.

[2]Center for Integrated Solutions for Infectious Diseases, Broad Institute, Cambridge, Massachusetts, United States of America.

*181 Longwood Ave, Boston, Massachusetts 02115, USA; (BEG); (YS).

[#]These authors contributed equally.

**Supplementary Figure Legends**

**Fig. S1**

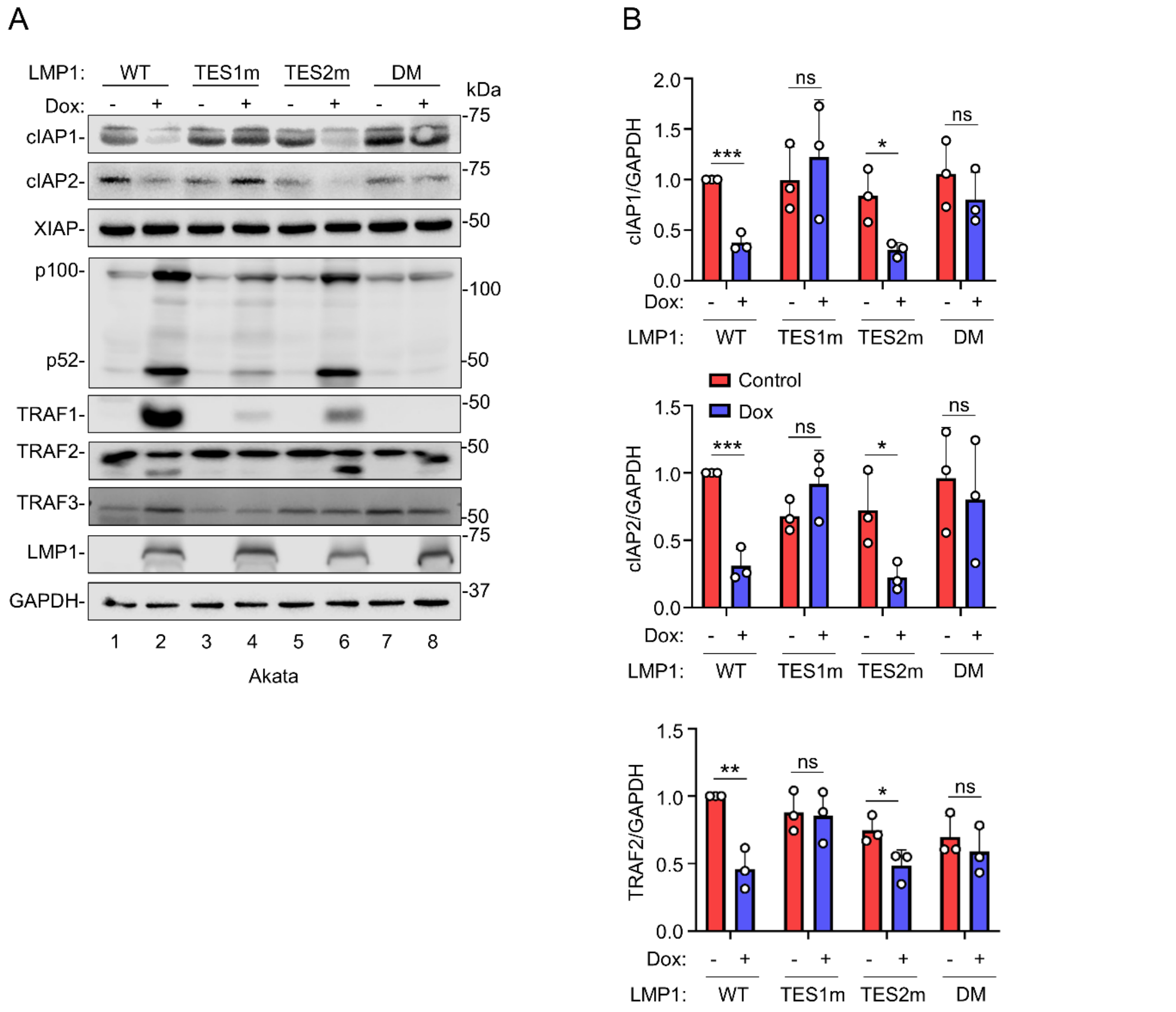


**Figure S1. LMP1 TES1 signaling downregulates cIAP1/2 and TRAF2 in Akata B cells.**

(A) Analysis of LMP1 TES1 vs TES2 signaling effects on cIAP1/2 and TRAF levels. Immunoblot analysis of WCL from Akata cells induced for WT, TES1m, TES2m or DM LMP1 expression by 250 ng/mL Dox for 24 hours. Blots are representative of n=3 experiments.

(B) Relative fold changes + SD of GAPDH load-controlled cIAP1, cIAP2 or TRAF2 values, based on densitometry from n=3 replicates of immunoblots as shown in (A). Values in vehicle control treated cells uninduced for WT LMP1 expression were set to 1.

Statistical significance was assessed by two-tailed unpaired Student’s t-test (B). ns, not significant, *p<0.05, **p<0.01, ***p<0.001.

**Fig. S2**

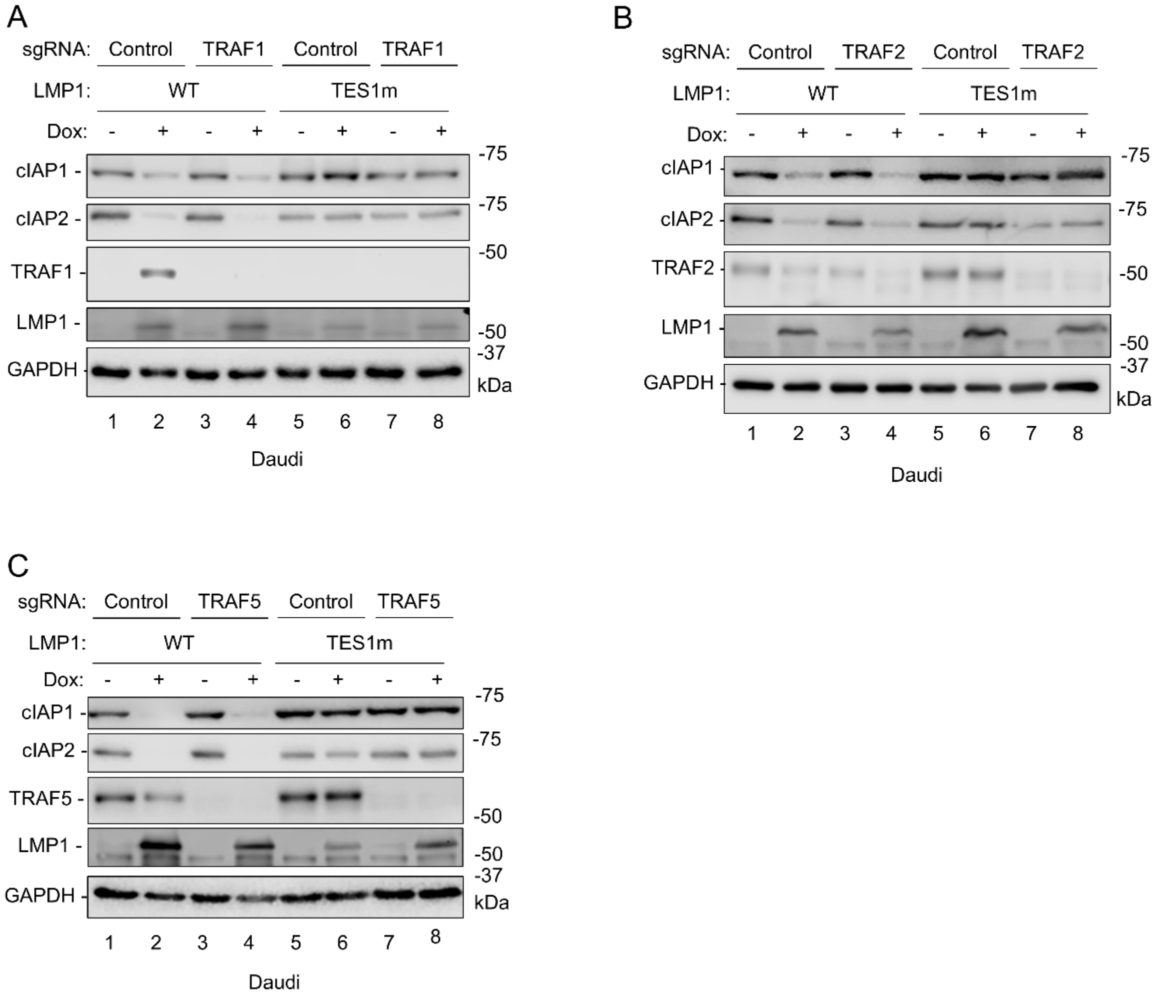


**Figure S2. TRAFs1, 2 and 5 are dispensable for LMP1 induced-cIAP1/2 depletion.**

(A) Immunoblot analysis of WCL from Cas9+ Daudi cells that expressed control or TRAF1 targeting sgRNA and that were induced for LMP1 WT or TES1m expression by 250ng/mL Dox for 24 hours.

(B) Immunoblot analysis of WCL from Cas9+ Daudi cells that expressed control or TRAF2 targeting sgRNA and that were induced for LMP1 WT or TES1m expression by 250ng/mL Dox for 24 hours.

(C) Immunoblot analysis of WCL from Cas9+ Daudi cells that expressed control or TRAF5 targeting sgRNA and that were induced for LMP1 WT or TES1m expression by 250ng/mL Dox for 24 hours.

Blots are representative of n=3 experiments.

Fig. S3

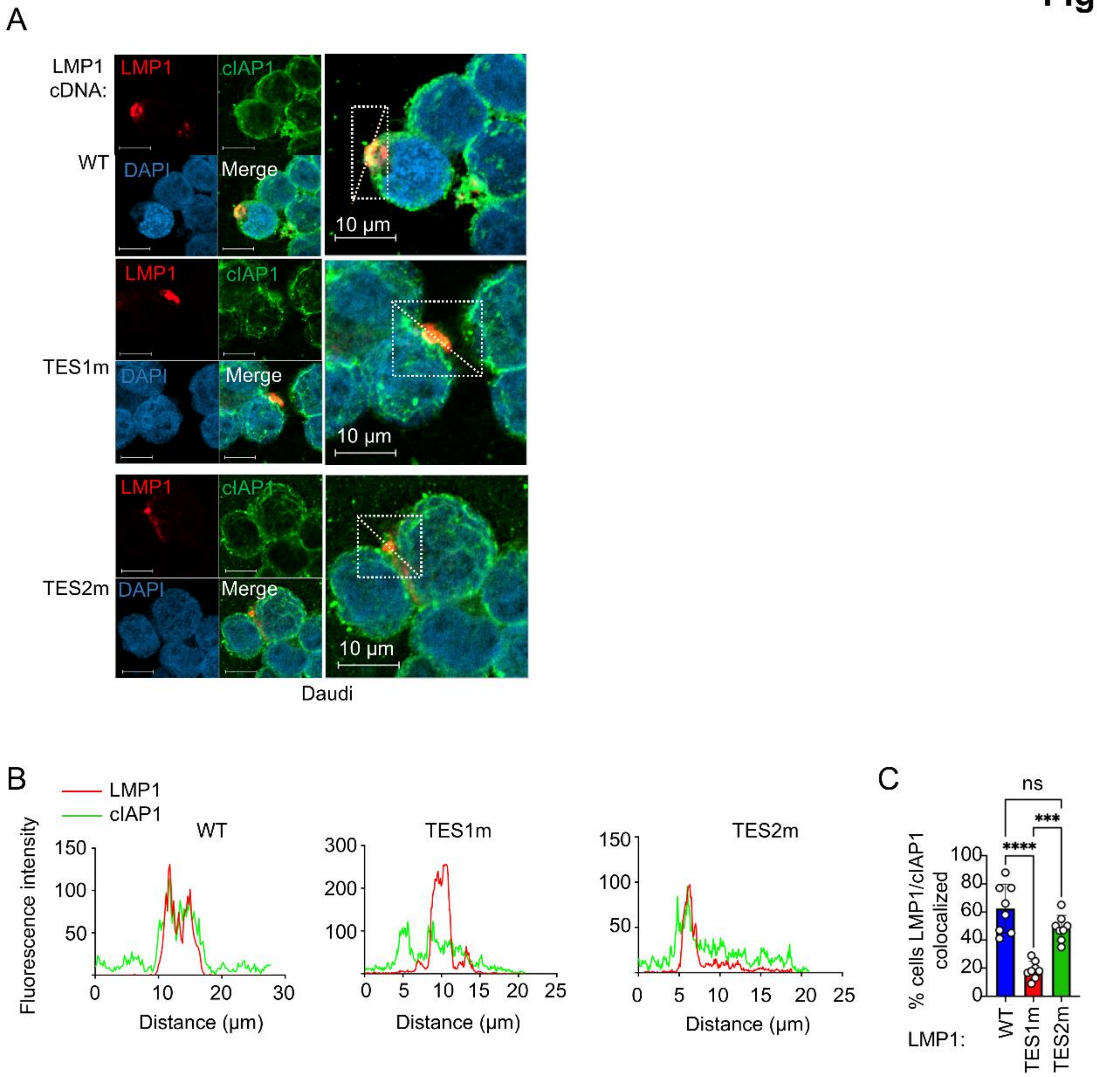


**Figure S3. LMP1 co-localizes with cIAP1 in a TES1 domain-dependent manner**

(A) Immunofluorescence analyses of cIAP1 and LMP1 localization in Daudi cells. WT, TES1m, TES2m or DM LMP1 expression was induced by 250ng/mL Dox for 24 hours, followed by treatment with 5 μM MG132 for 6 hours. Images are representative of 10 randomly chosen fields per sample.

(B) Line scanning of cIAP1 (green) and LMP1 (red) fluorescence intensity within the annotated the white rectangles shown in panel A.

(C) Quantification of cells with overlapping cIAP1 and LMP1 signal in Daudi cells. Line scanning was performed with Zeiss Zen Lite (Blue) software. Statistical significance was assessed by one-way ANOVA followed by Tukey's multiple comparisons test. ns, not significant, ***p<0.001, ****p<0.0001.

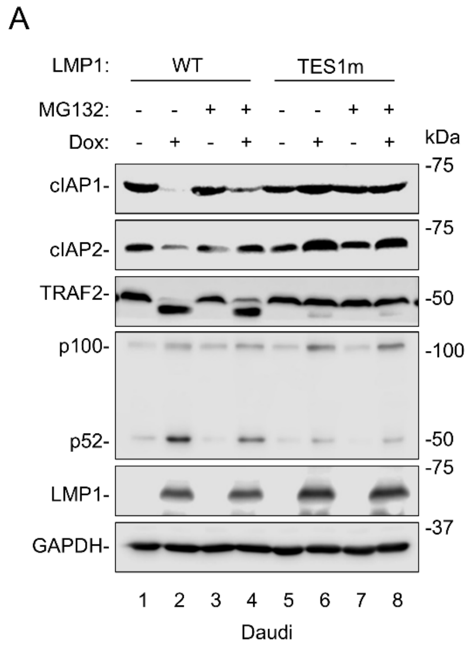


**Figure S4. MG132 partially rescued LMP1-induced degradation of TRAF2.**

Immunoblot analysis of WCL from Daudi cells induced for WT LMP1 expression by 250ng/mL Dox for 24 h, followed by treatment with 5μM MG132 for 8 hours. Blots are representative of n=3 experiments.
